# Estimating parameters from multiple time series of population dynamics using Bayesian inference

**DOI:** 10.1101/392449

**Authors:** Benjamin Rosenbaum, Michael Raatz, Guntram Weithoff, Gregor F. Fussmann, Ursula Gaedke

**Author notes:** Benjamin Rosenbaum and Michael Raatz contributed equally to this work.

## Abstract

1. Empirical time series of interacting entities, e.g. species abundances, are highly useful to study ecological mechanisms. Mathematical models are valuable tools to further elucidate those mechanisms and underlying processes. However, obtaining an agreement between model predictions and experimental observations remains a demanding task. As models always abstract from reality one parameter often summarizes several properties. Parameter measurements are performed in additional experiments independent of the ones delivering the time series. Transferring these parameter values to different settings may result in incorrect parametrizations. On top of that, the properties of organisms and thus the respective parameter values may vary considerably. These issues limit the use of a priori model parametrizations.
2. In this study, we present a method suited for a direct estimation of model parameters and their variability from experimental time series data. We combine numerical simulations of a continuous-time dynamical population model with Bayesian inference, using a hierarchical framework that allows for variability of individual parameters. The method is applied to a comprehensive set of time series from a laboratory predator-prey system that features both steady states and cyclic population dynamics.
3. Our model predictions are able to reproduce both steady states and cyclic dynamics of the data. Additionally to the direct estimates of the parameter values, the Bayesian approach also provides their uncertainties. We found that fitting cyclic population dynamics, which contain more information on the process rates than steady states, yields more precise parameter estimates. We detected significant variability among parameters of different time series and identified the variation in the maximum growth rate of the prey as a source for the transition from steady states to cyclic dynamics.
4. By lending more flexibility to the model, our approach facilitates parametrizations and shows more easily which patterns in time series can be explained also by simple models. Applying Bayesian inference and dynamical population models in conjunction may help to quantify the profound variability in organismal properties in nature.

## Introduction

Trophic interactions provide the elementary link in all food webs. Long-term data sets of such interactions become increasingly available, both from field observations, and highly controlled large-scale and laboratory experiments (G. F. Fussmann et al. 2000; Tirok and Gaedke 2007; Becks et al. 2010; Magurran et al. 2010; Weigelt et al. 2010; Becks et al. 2012; Boit and Gaedke 2014). To mechanistically understand these trophic interactions, models are employed with the goal of reproducing the observed population dynamics, which can either settle to a steady state, or include more complex patterns like limit cycles or chaos (G. F. Fussmann et al. 2000; Boit et al. 2012; Becks and Arndt 2013; Barraquand et al. 2017). Obtaining an agreement of such non-static experimental observations and modelled time series remains a demanding task. Often, modellers are faced with situations where they can reproduce basic features of the dynamics, but not on the right time scales or not on biomass levels close to the data. One logical reaction would be to refine the model structure and include a higher level of biological detail. However, another reason for such disagreements between model and data may be a generally valid model structure but an incorrect parametrization. In this study, we present a method to obtain the relevant model parameters directly from the experimental data by applying Bayesian inference to a comprehensive set of time series data for a laboratory predator-prey system that features both steady states and cyclic population dynamics.

Incorrect parametrizations can arise for different reasons. Firstly, a model always has to abstract from reality. This implies that individual model parameters summarize a multitude of different ecological processes. The impact of these processes on the model parameter likely changes over time. Due to these non-modelled processes a one-to-one relationship between empirically determined parameter values and model parameters is impossible. For example, typical predator-prey models consider a conversion efficiency by which prey biomass is converted into predator growth. Among others this efficiency depends on the variable prey abundance (S. Schälicke *in prep*.) and the relative importance of basal and activity respiration (Kath et al. 2018). Secondly, model parameters are often obtained experimentally under slightly different settings than the actually observed population dynamics. An example would be to measure the prey’s population growth rate in batch experiments and then use this parameter in a chemostat model. By design, growth conditions in batch and chemostat cultures differ in some aspects, such as the dynamics of nutrient limitation or the selection pressure, e.g. for higher nutrient affinities versus maximum growth rates. Therefore, the parameters that were obtained for one experimental setting might be of limited value for a different one.

It is more and more recognized that the functional traits of organisms that determine trophic interactions comprise a considerable variability (Litchman and Klausmeier 2008; Bolnick et al. 2011; Violle et al. 2012; Bolius et al. 2017; Gaedke and Klauschies 2017). This trait variability can have far-reaching consequences both at the population level (Abrams 1999; Post and Palkovacs 2009; Becks et al. 2010; Ehrlich et al. 2017; Raatz et al. 2017; Cortez 2018) and at the community level (McGill et al. 2006; Hillebrand and Matthiessen 2009). In such cases, employing just one parametrization for different time series of the same system may be insufficient and hamper the agreement of experimental and modelled population dynamics. Instead, the model will potentially support parts of the data or comprise certain general features, but will fail to reproduce its entire behaviour. Our Bayesian framework allows to retrieve information on such parameter-related uncertainties directly from the time series data, which might even become apparent as between-replicate differences, and provide individual parameter estimates for each data set.

Dynamical population models have traditionally been fitted to time series data by least squares or maximum likelihood methods (Costantino et al. 2005; Cao et al. 2008; DeLong et al. 2014; Rall and Latz 2016; Curtsdotter et al. 2018), see Bolker (2008) and Aster et al. (2012) for a general introduction. While they offer point estimates for unknown parameters, their confidence intervals rely on a local approximation and a normality assumption of the likelihood function (Bolker 2008, Chap. 6.5).

Bayesian methods, on the other hand, quantify uncertainty more precisely by globally exploring the parameters’ posterior probability distribution using Markov chain Monte Carlo (MCMC) sampling. They allow, for instance, direct inference on sought parameters and derived quantities, utilizing prior information, defining hierachical levels among parameters, and recovering unobserved system states (Kindsvater et al. 2018).

For discrete-time population dynamics, Bayesian methods have received growing attention over the last years (Almaraz and Oro 2011; Elderd and Miller 2015; Wittwer et al. 2015; Compagnoni et al. 2016; Robinson et al. 2017). In a discrete setup, state-space models (SSM) are feasible and allow, e.g., the separation of process and observation error (Hefley et al. 2013), recovering latent states (Hosack et al. 2012), incorporating age-structure (Taboadai and Anadón 2016), adding environmental covariates (Almaraz et al. 2012; Koons et al. 2015), or spatially explicit models (Iijima et al. 2013). These advances were facilitated by the probabilistic programming environments BUGS (Lunn et al. 2009) and JAGS (Plummer 2003).

The implementation of continuous-time population dynamics (described by ordinary differential equations, ODEs) is available in BUGS but not in JAGS. Until recently, modellers often combined numerical simulations of ODEs manually with MCMC routines (Gilioli et al. 2008; Toni et al. 2009; Johnson et al. 2013; Smith et al. 2015; Papanikolaou et al. 2016; Boersch-Supan et al. 2017). The probabilistic programming language Stan (Carpenter et al. 2017) offers an integrated solution. It comes with interfaces to R, Python, Matlab and more, a built-in numerical ODE solver and a Hamiltonian Monte Carlo (HMC) sampler (Monnahan et al. 2017). Thus, it supports fitting dynamical population models to time series data in a Bayesian framework, see K. E. Fussmann et al. (2017) and Carpenter (2018) for recent applications.

In this study, we will apply Bayesian inference in Stan to a set of time series of a predator-prey system in a chemostat, i.e. a continuous flow-through culture (Novick and Szilard 1950). The parameters of a well-established continuous-time chemostat ODE model will be estimated yielding posterior distributions for the parameters, which allow also to quantify their uncertainties. By comparing the posteriors for the individual time series we can deduce a variability among them that manifests in different types of population dynamics and pin-point to specific parameters that seem to determine this variability.

## Materials and Methods

### Data collection

Chemostat experiments were performed to obtain predator-prey time series at a high temporal resolution in a highly controlled environment (Guntram Weithoff, Bernd Blasius, Gregor F. Fussmann, Ursula Gaedke, *et al*., unpublished) resulting in a large collection of long-term time series with several different species. From these we selected a subset of 13 experiments, which were replicates in the sense that the same species were used at the same inflow nutrient concentration and dilution rate, and daily counts of prey and predators are available. The experiments were conducted with a metazoan predator, the rotifer *Brachionus calyciflorus* s.s. (Michaloudi et al. 2018; Paraskevopoulou et al. 2018), and its prey *Monoraphidium minutum*, a unicellular green alga. The algae grew on nitrogen-limited medium. However, daily nitrogen concentrations are not available. The experiments were performed within a time span of seven years, with some individual experiments lasting longer than one year. They yielded time series which differed with respect to the degree that they exhibited more or less regular cyclic dynamics, or more or less constant predator and prey densities. From these 13 replicates we selected all shorter time series that showed either clear and pronounced predator-prey cycles or steady-state equilibria, and excluded pronounced initial transient phases. We chose a minimum sample length of 20 days which would typically allow for at least two predator-prey cycles in this system. This process resulted in a set of 18 samples, 10 of which featured a steady state and eight contained cyclic dynamics.

### Dynamical population model

The continuous-time population dynamics of a predator-prey system in a chemostat with nitrogen *S*, algae *A* and rotifers *R* are described by the equations:

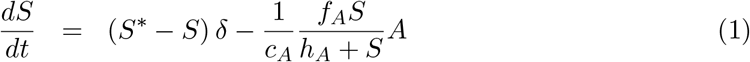

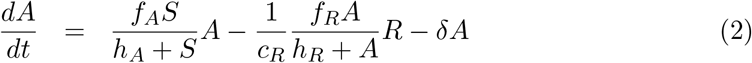

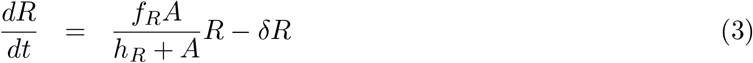

This formulation represents a slightly simplified version of the one originally presented by G. F. Fussmann et al. (2000) and neglects age structure of the predator. *δ* is the system’s inflow and outflow rate, and the concentration of nutrients in the inflow is given by *S*\*. The factors *c_A_* and *c_R_* define the conversion of nutrients into algal biomass and algal into rotifer biomass, respectively. The growth rate of algae is described by Monod kinetics 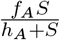. *f_A_* is the maximum growth rate and *h_A_* is the half-saturation constant (the resource density at which the growth rate equals half of the maximum growth rate). The same applies to the resource-dependent growth rate of rotifers feeding on algae 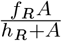, which is described by a type II functional response (Real 1977). The maximum growth rates *f_A_*[d^−1^], *f_R_*[d^−1^] and the conversion factors *c_A_*[cells μmol^−1^], *c_R_*[ind cells^−1^] are free parameters, which are estimated as described in the next section. We used constant values for the following parameters from a similar system which instead used *Chlorella vulgaris* as the prey species: *S*\* = 80 μmol l^−1^ and *δ* = 0.55 d^−1^, as they were carefully controlled in the experiments, and *h_A_* = 4.3 μmol l^−1^ and *h_R_* = 7.5 · 10^8^ cells l^−1^ (G. F. Fussmann et al. 2000). These values for the half-saturation constants were in the range of our predicted and observed resource states *S* and *A*, respectively (cf. Figs. 2,3). We chose to not use half-saturation constants *h_A_* (or *h_R_*) as free parameters, since the estimates can be highly correlated with maximum growth rates *f_A_* (or *f_R_*). Their combined effects on the resource-dependent growth rate 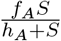 (or 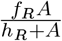) can only be disentangled if the data cover a large range in the resource states *S* (or *A*) (Rosenbaum and Rall 2018), which is not the case for the present chemostat experiments.

### Model fitting and inference

We combined numerical simulations of the deterministic dynamical population model (eqns. 1–3) with Bayesian parameter estimation by drawing samples from the posterior probability distribution *P*(*θ*|*y*) of the free parameters *θ* given the data *y*, based on the likelihood *P*(*y*|*θ*) and the prior distribution *P*(*θ*). We used Hamiltonian Monte Carlo sampling in Stan, accessed via the `RStan R`-package (Stan Development Team 2018). The Stan software comes with a built-in Runge-Kutta ODE solver with adaptive stepsize control for generating predictions *ŷ*(*θ*).

The likelihood calculation *P*(*y*|*θ*) is carried out automatically by the software when provided with predictions *ŷ*(*θ*) and the distribution of residuals *ŷ*(*θ*) — *y*. The predictions *ŷ*(*θ*) are defined by the numerical solutions of the ODE *Â*(*t_i_*) and *Ȓ*(*t_i_*), evaluated at times *t_i_*, for a given parameter combination *θ*. We chose a log-normal distribution of the residuals, i.e. 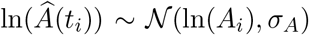 and 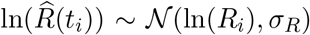, with scale parameters *σ_A_* and *σ_R_*. This trajectory matching method technically corresponds to treating the model deterministically and to assuming pure observation errors in the data without any process error (Bolker 2008, Chap. 11). Note that, even without data for the concentration nitrogen *S*, it is possible to fit the ODE model by including the initial densities of the predictions 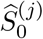, 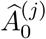 and 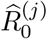 as free parameters (Carpenter 2018). However, one-step-ahead fitting (i.e., assuming pure process error) would only be possible for this ODE model if data for all state variables *S*, *A* and *R* was available. We did not consider full state-space models accounting for both process and observation error.

We fitted the maximum growth rates *f_A_* and *f_R_* and the conversion factors *c_A_* and *c_R_* on their logarithmic scale (see model code, Supporting Information). The dynamics and the statistical model are equivalent to fitting them on their original scale. But since they differ by several orders of magnitude, we found that log-transforming the parameter search space makes the iterative MCMC routine more robust.

We used a *hierarchical model* for the maximum growth rates *f_A_* and *f_R_* and for the conversion factors *c_A_* and *c_R_* by using time series identity *j* = 1,…, *m* as a *random effect*. This means that every time series *j* is fitted with its individual set of parameters 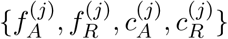, while each of these four parameters originates from a joint distribution across all *m* time series replicates. Thus, some information is shared across the individual replicates via the joint distribution, therefore this technique is also known as *partial pooling*. In a Bayesian framework, this can be modeled via hierarchical dependencies in the prior distributions. Including logarithmic scaling, they read

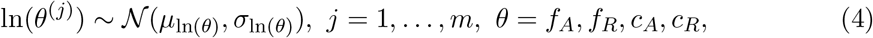

where *μ*_ln(*θ*)_ are the overall means and *σ*_ln(*θ*)_ the standard deviations across all *m* time series. *μ*_ln(*θ*)_ and *σ*_ln(*θ*)_ are also free parameters with their own prior distribution (see Table 1 for a full description of the priors).

**Table 1:** Free parameters and their prior distributions. 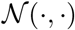 denotes a normal distribution, 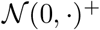 denotes a positive half-normal distribution truncated at zero.

| parameter |  | distribution | type |
| --- | --- | --- | --- |
| max. growth rates | $\ln(f_A^{(j)})$ | $\mathcal{N}(\mu_{\ln(f_A)}, \sigma_{\ln(f_A)})$ | hierarchical |
| max. growth rates | $\ln(f_R^{(j)})$ | $\mathcal{N}(\mu_{\ln(f_R)}, \sigma_{\ln(f_R)})$ | hierarchical |
| conversion factors | $\ln(c_A^{(j)})$ | $\mathcal{N}(\mu_{\ln(c_A)}, \sigma_{\ln(c_A)})$ | hierarchical |
| conversion factors | $\ln(c_R^{(j)})$ | $\mathcal{N}(\mu_{\ln(c_R)}, \sigma_{\ln(c_R)})$ | hierarchical |
| overall mean | $\mu_{\ln(f_A)}$ | $\mathcal{N}(1.194, 1)$ | weakly informative |
| overall mean | $\mu_{\ln(f_R)}$ | $\mathcal{N}(0.811, 1)$ | weakly informative |
| overall mean | $\mu_{\ln(c_A)}$ | $\mathcal{N}(17.728, 1)$ | weakly informative |
| overall mean | $\mu_{\ln(c_R)}$ | $\mathcal{N}(-10.597, 1)$ | weakly informative |
| overall standard deviation | $\sigma_{\ln(f_A)}$ | $\mathcal{N}(0, 1)^+$ | weakly informative |
| overall standard deviation | $\sigma_{\ln(f_R)}$ | $\mathcal{N}(0, 1)^+$ | weakly informative |
| overall standard deviation | $\sigma_{\ln(c_A)}$ | $\mathcal{N}(0, 1)^+$ | weakly informative |
| overall standard deviation | $\sigma_{\ln(c_R)}$ | $\mathcal{N}(0, 1)^+$ | weakly informative |
| initial densities | $\ln(\hat{S}_0^{(j)})$ | $\mathcal{N}(2, 2)$ | weakly informative |
| initial densities | $\ln(\hat{A}_0^{(j)})$ | $\mathcal{N}(20, 2)$ | weakly informative |
| initial densities | $\ln(\hat{R}_0^{(j)})$ | $\mathcal{N}(10, 2)$ | weakly informative |
| residual standard deviation | $\sigma_A$ | $\mathcal{N}(0, 2)^+$ | weakly informative |
| residual standard deviation | $\sigma_R$ | $\mathcal{N}(0, 2)^+$ | weakly informative |

We tested for variation in the dynamics of the different time series by uncovering differences in parameters. For each parameter *θ* = *f_A_*, *f_R_*, *c_A_*, *c_R_*, pairwise contrasts *θ*^(*j*)^ – *θ*^(*k*)^ between two time series *j* and *k* were inferred. I.e., we generated posterior probabilities *P_jk_* = *P*(*θ*^(*j*)^ > *θ*^(*k*)^) that quantify these differences. These quantities *P_jk_* are directly computed from the posterior distribution by dividing the number of samples featuring *θ*^(*j*)^ > *θ*^(*k*)^ by the total number of samples.

To further investigate the importance of variation among the parameter estimates for different time series, we also fitted the ODE model (eqns. 1–3) using a single set of parameters {*f_A_,f_R_,c_A_,c_R_*} for all 18 time series as a null model. In contrast to the hierarchical (partial pooling) model above, this is also known as *complete pooling*, since all information across individual replicates is combined. Only for the initial states 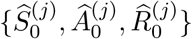 we allowed distinct values for each of the time series *j* = 1,…, 18.

We briefly comment on numerical issues that can arise when combining numerical solutions of ODEs with MCMC sampling. When the MCMC sampler explores the parameter space, points can be sampled that make the computation of the likelihood by numerical simulation of the ODE infeasible (e.g. by requiring an immensely small integration step-size or simply by divergent state variables). Still, the sampling algorithm requires the computation of the likelihood to proceed with the iterations. To prevent the sampler from entering regions of the parameter space where, over a whole range of values, the likelihood is not available, we used two strategies. First, we implemented a numerical condition which prevents the numerical ODE solution from diverging or crossing the lower boundary of zero by setting the right-hand-side of the ODE to zero if one of the state variables exceeds a reasonable range of [10^−6^, 10^16^] (see model code in Supporting Information). Second, we used weakly informative priors on the overall mean parameters *μ*_ln(*θ*)_ based on measured values from G. F. Fussmann et al. (2000): 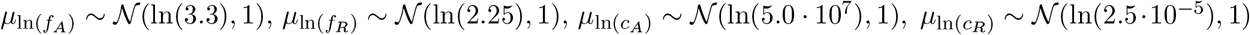 (see also Table 1). We used the same priors for the *complete pooling* model.

### Simulation study

Before fitting the presented models to the experimental chemostat data, we first validated our modeling approach in a simulation study. By fitting the hierarchical model to simulated time series, which were generated by known parameters, we tested the identifiability of model parameters. These parameters 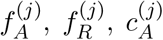 and 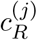 were drawn randomly from lognormal distributions using the measured values from above (G. F. Fussmann et al. 2000) as means and a standard deviation of 0.5. Initial states of the time series were also assigned randomly according to Table 1. We numerically simulated ODE trajectories of eqns. 1–3 for 100 days and chose 10 time series that settled to different steady states and 10 time series that featured cyclic dynamics of different frequencies and amplitudes. We used the observations of algal and rotifer states of the last 20 days (leaving out nitrogen states as in the experimental data) and added a random error with zero mean and standard deviation of 0.1 on the ln-scale (see also Figs. A1,A2, Supporting Information).

## Results

### Model convergence

We fitted all models (hierarchical model in simulation study; hierarchical model and complete pooling model for experimental chemostat data) by running 10 individual MCMC chains in parallel with an adaptation phase of 1,000 iterations and a sampling phase of 5,000 samples each, summing up to 50,000 samples of the posterior distribution. The runtime was approximately 7 days on a 2.2GHz Intel Xeon server architecture. Visual inspection of the trace plots and density plots showed a good mixture of the chains. Gelman-Rubin statistics of *Ȓ* < 1.01 and an adequate effective sample size *n*_eff_ (i.e. the estimated number of independent samples) verified convergence (Gelman and Hill 2007). See Supporting Information (Tables A1–A4) for a full list of parameter estimates and their statistics.

### Identifiability of parameters

We used the simulation study to assess if known parameters can be recovered accurately by fitting the hierarchical model to a synthetic set of 10 steady-states and 10 cyclic time series (cf. Figs. A1,A2, Supporting Information). Fig. 1 shows the posterior error distributions (distributions of estimated parameter values minus true values on ln-scale) of maximum growth rates 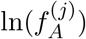 and 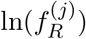 and conversion factors, 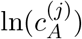 and 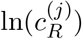 of prey and predators, respectively. We found that all parameters of cyclic time series 11–20 were accurately identifiable. The posterior medians generally did not deviate more than 0.05 from the true parameters and all estimates featured a low uncertainty (posterior standard deviations smaller than 0.04).

**Figure 1:**
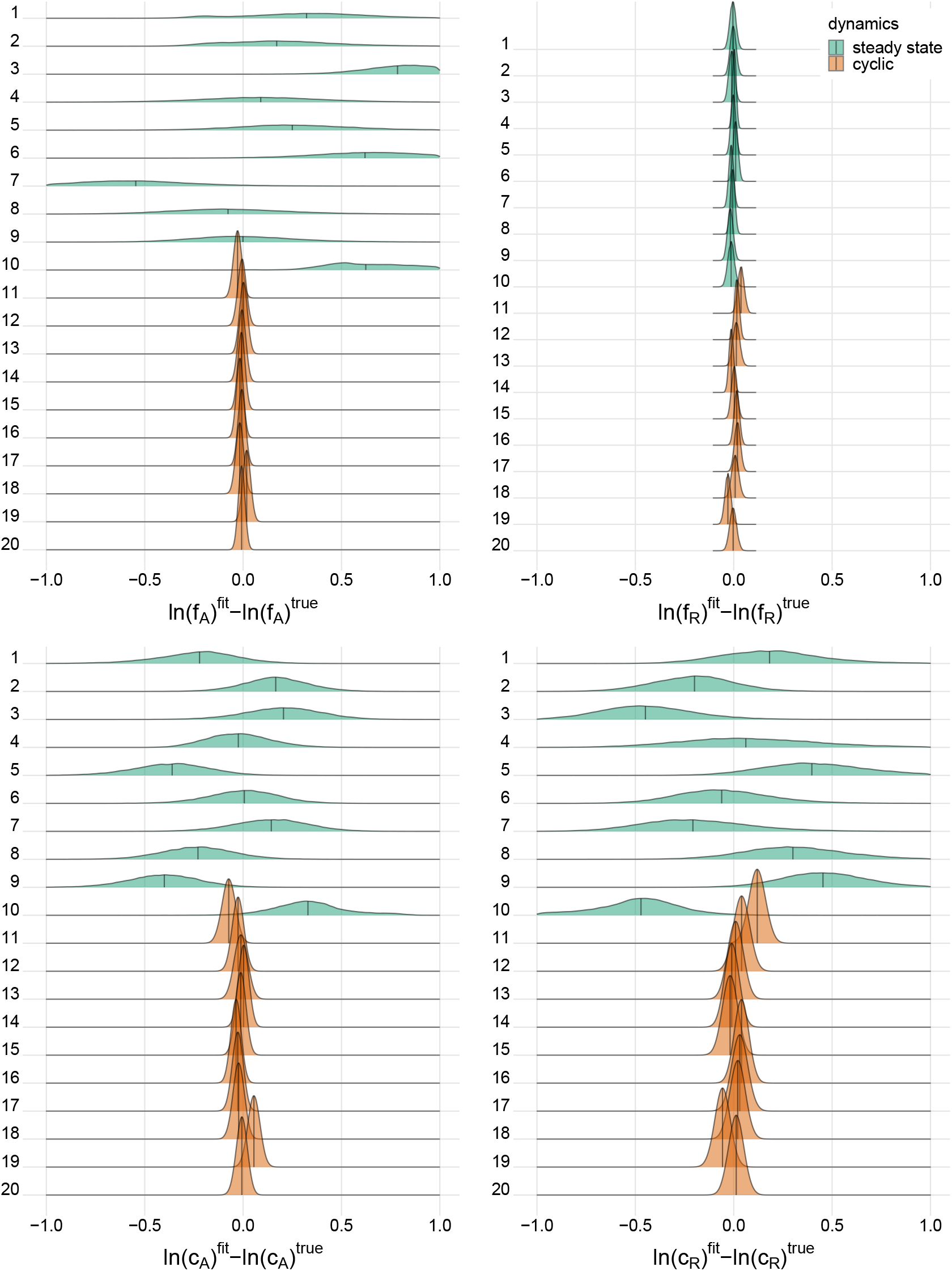
**Posterior error distributions** of maximum growth rates 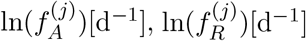, and conversion factors 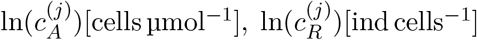 for fitting **simulated time series** (*j* = 1,…, 20). Simulated trajectories 1–10 featured a steady-state equilibrium (green), while simulated trajectories 11–20 featured cyclic behavior (orange). Vertical bars indicate median.

For steady-state time series 1–10, however, posterior distributions of algal maximum growth rates 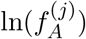 showed relatively high uncertainties (posterior medians deviating up to 0.8 from true values with posterior standard deviations up to 0.34). We assume that, in combination with the lack of nitrogen data *S*, steady-state time series, which cover a smaller range in the state space, contain less information about the resource density-dependent growth rates of algae feeding on nitrogen 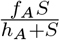 than cyclic time series. Estimates for rotifer maximum growth rates 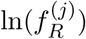, on the other hand, were highly accurate just as for cyclic time series. In contrast to *f_A_*, the data seem to provide enough information on the growth rates of rotifers feeding on algae 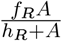 even in steady-state time series, since observations for both involved trophic levels are available. The estimates for conversion factors 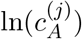 and 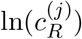 were also less accurate for steady-state time series 1–10 than for cyclic time series 11–20 (posterior medians deviate up to 0.47 from true parameters, with an uncertainty of posterior standard deviations up to 0.37). We assume that steady-state data is not as informative as cyclic data on the conversion factors, as we also observe a high correlation between the posterior of 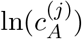 and 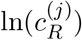 for every steady-state time series (cf. Supporting Information, Fig. A3). Note that correlation per se does not imply unidentifiability. We observed correlations between the posteriors of cyclic time series parameters as well (cf. Supporting Information, Fig. A4), while these parameters are estimated with a high accuracy.

### Posterior predictions

After assessing the identifiability of the hierarchical model in a simulation study, we investigated its performance with the experimental chemostat data. We generated posterior predictions by numerical simulations of the ODE system (eqns. 1–3) with all samples of the posterior distribution (Figs. 2,3). After a short transient phase, the median predicted trajectories feature either a steady-state equilibrium (time series 1–9), cyclic behavior (time series 13–18); or the posterior distribution includes parameter samples producing steady states as well as cyclic trajectories (time series 10–12). Correspondingly, we found multimodalities in the posterior distributions of time series 10–12 (see Supporting Information, Figs. A6&A7).

**Figure 2:**
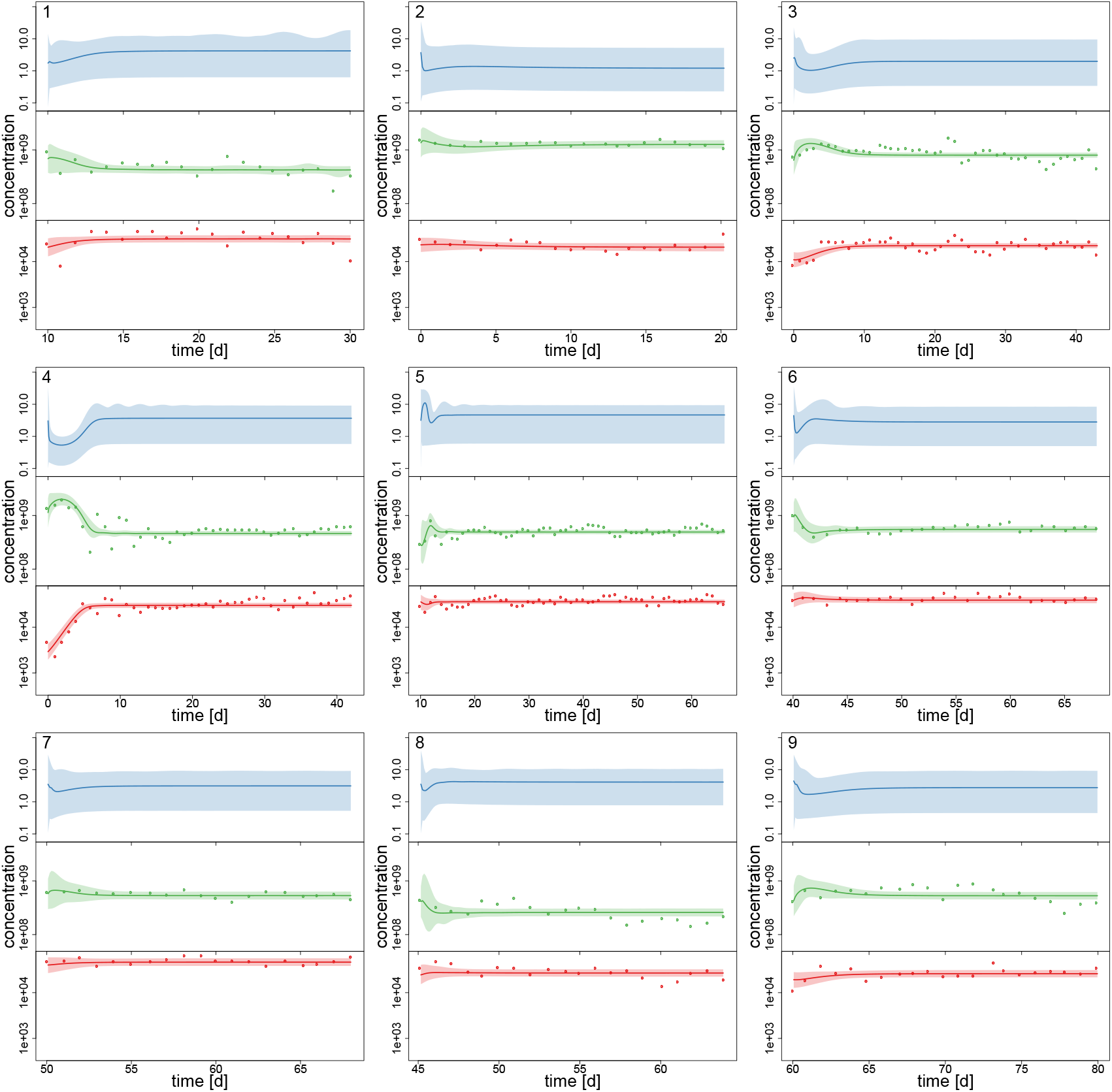
**Experimental time series** 1–9 of 18, data (dots) and posterior predictions of hierarchical model for nitrogen [μmol l^−1^] (blue), algae [cells l^−1^] (green) and rotifers [ind l^−1^] (red). Solid lines represent median predictions, shaded areas depict 95% highest density intervals of the predictions. Predicted trajectories 1–9 featured a steady-state equilibrium.

**Figure 3:**
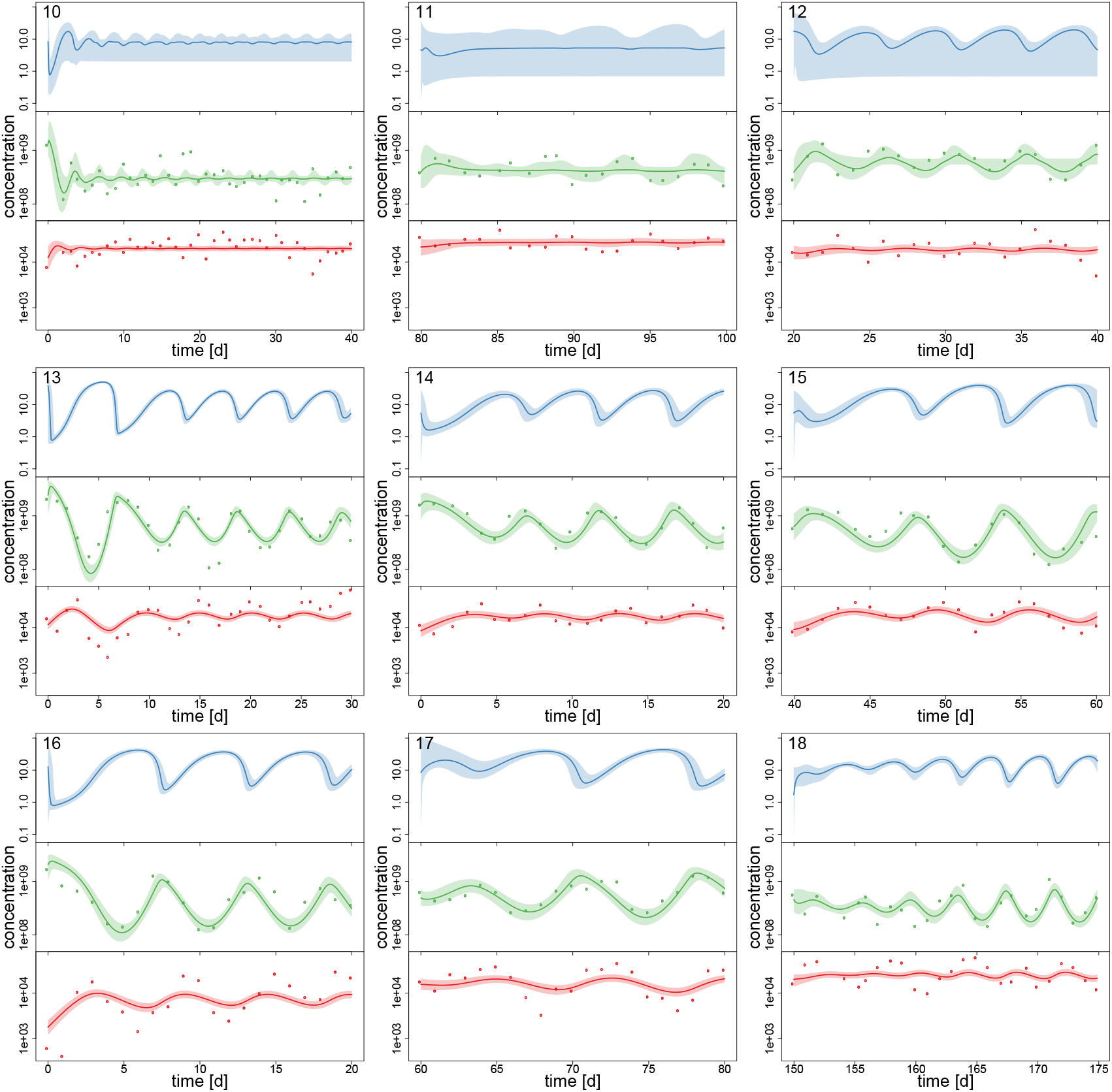
**Experimental time series** 10–18 of 18, data (dots) and posterior predictions of fitted model for nitrogen [μmol l^−1^] (blue), algae [cells l^−1^] (green) and rotifers [ind l^−1^] (red). Solid lines represent median predictions, shaded areas depict 95% highest density intervals of the predictions. Predicted trajectories 13–18 featured cyclic behavior. Time series 10–12 featured multimodalities in the posterior distribution and the predictions did not exhibit a clear tendency towards a steady state or cycles.

Interestingly, the relative uncertainty of the predictions (quantified by 95% confidence intervals) for all state variables is substantially reduced in time series 13–18 where the predictions feature cyclic behavior compared to other time series (Figs. 2,3). We measured the predictive accuracy in the univariate time series by normalized root mean-square-errors (cf. Fig. A7, Supporting Information). Here we see that the error distributions are shifted to smaller values and become more narrow for cyclic dynamics, also indicating a better fit. Again, this may be explained by acknowledging that steady-state data, which covers a smaller range in the state space than cyclic data, contains less information about the process rates and hence the parameters.

As no data constrains the predictions for nitrogen, the uncertainty is even higher here than in algae or rotifers in time series 1–9 (this was also observed for steady-state time series in the simulation study, cf. Supporting Information Fig. A1). Also, we found that our ODE model is able to predict the full amplitude of cycles in algal states better than in rotifer states (Figs. 2,3; Fig. A7, Supporting Information). This is likely caused by a higher regularity in the algal data, which covers a larger amplitude decreasing the relative counting error. Also, algae feature a less complex life cycle than rotifers and their dynamics should thus be less variable.

### Variation among time series

For assessing the variation in the parameters across the experimental chemostat time series (*j* = 1,…, 18), we plotted the marginal (i.e. one-dimensional projections of the multivariate) posterior probability distributions of the logarithmic maximum growth rates 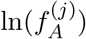 and 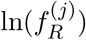 and the logarithmic conversion factors, 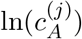 and 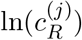 of prey and predators, respectively (Fig. 4). We also computed probabilities of pairwise contrasts *P_jk_* = *P*(*θ*^(*j*)^ > *θ*^(*k*)^) for a more detailed examination of the differences across time series (*θ* = *f_A_*, *f_R_*, *c_A_*, *c_R_*, Tables 2–5). Values close to one or close to zero indicate significant pairwise differences. Note that the tables are symmetrical in the sense of *P_jk_* = 1 – *P_kj_*.

**Figure 4:**
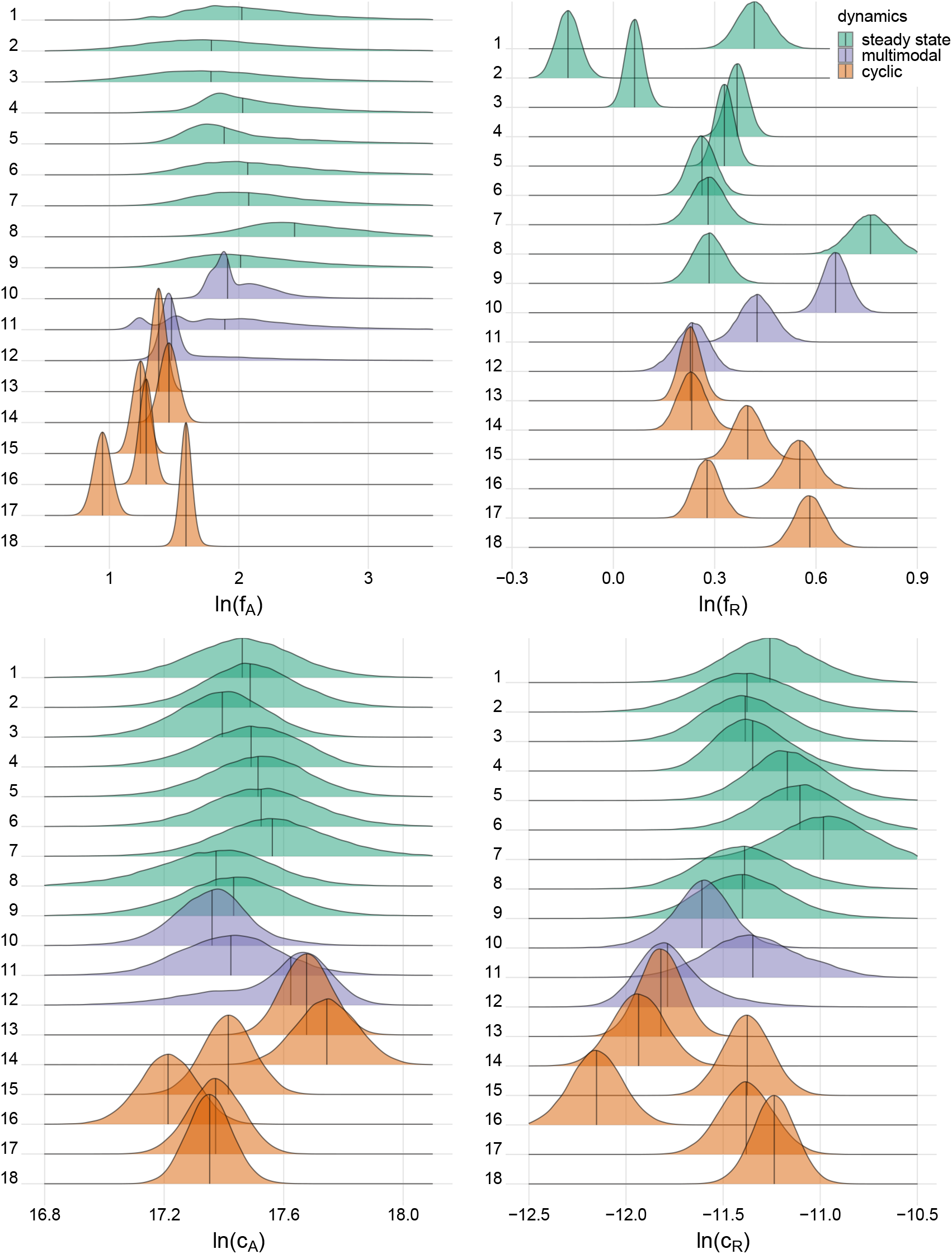
Marginal **posterior distributions** of maximum growth rates 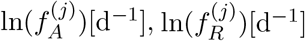, and conversion factors 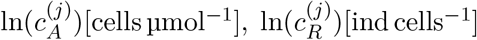 for fitting **experimental time series** (*j* = 1,…, 18). Predicted trajectories 1–9 featured a steady-state equilibrium (green), while predicted trajectories 13–18 featured cyclic behavior (orange). Time series 10–12 (purple) featured multimodalities in the posterior distribution and the predictions did not exhibit a clear tendency towards a steady state or cycles. Vertical bars indicate median.

We found that time series with predicted steady states feature systematically higher values of *f_A_* than time series with cyclic dynamics (Fig. 4a, Table 2, top right and bottom left blocks). While in cyclic time series 13–18 values of *f_A_* differ among each other (bottom right block), no evidence was found for differences among steady-state time series 1–9 (top left block). Steady-state time series also exhibit a high uncertainty in *f_A_* estimates with confidence intervals spanning from 2.43 d^−1^ to 40.85 d^−1^ when transformed back to a linear scale (Fig. 4a, see also Supporting Information, Table A1). This uncertainty is substantially reduced in cyclic time series 13–18 with confidence intervals spanning from 2.29 d^−1^ to 5.31 d^−1^. Here, the predicted parameter values are close to the published value in G. F. Fussmann et al. (2000), which however was published for *Chlorella* instead of *Monoraphidium*, but should be in the same range.

**Table 2:**
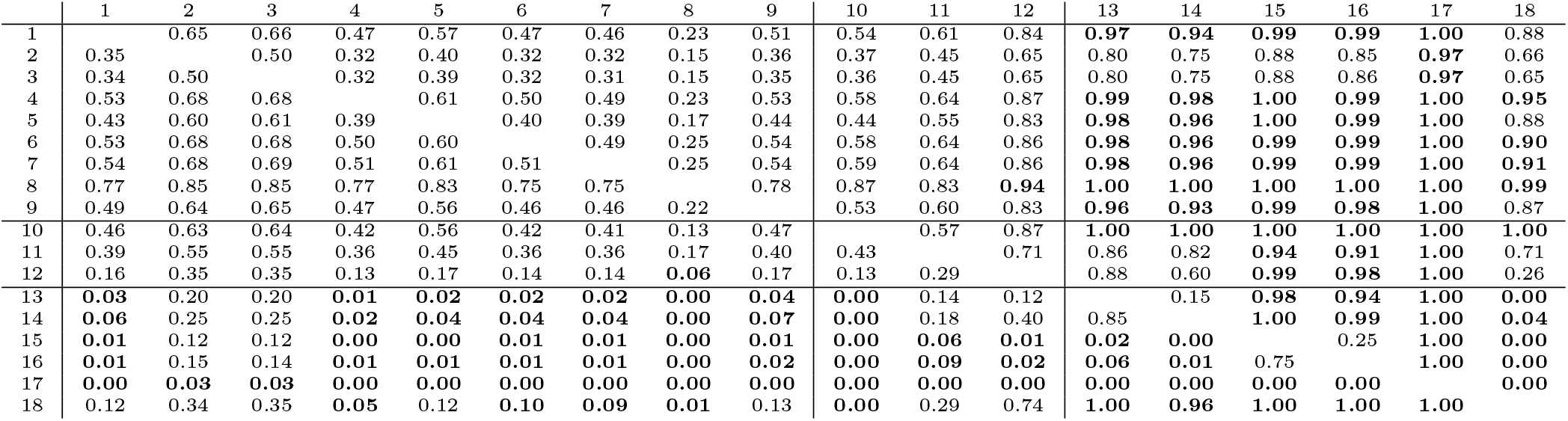
Pairwise contrasts 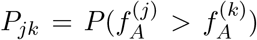 of maximum growth rates *f_A_*. Bold numbers indicate significant differences, quantified by probabilities *P_jk_* > 0.90 (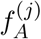 significantly larger than 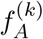) and probabilities 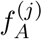 significantly smaller than 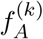). Block partitioning refers to time series 1–9 featuring a steady-state equilibrium, 10–12 having multimodal posterior distributions featuring both steady states and cycles, and 13–18 featuring cyclic dynamics.

For the rates *f_R_* we found pairwise differences across all time series (Fig. 4b, Table 3). No systematic effect of cyclic or steady-state time series was observed (i.e., *f_R_* estimates for cyclic time series are not systematically smaller than estimates for steady-state time series or vice versa). In contrast to the rates *f_A_*, even steady-state data provide enough information on rates *f_R_* (with consumer and resource data available), resulting in a low uncertainty in estimation just as for cyclic time series.

**Table 3:**
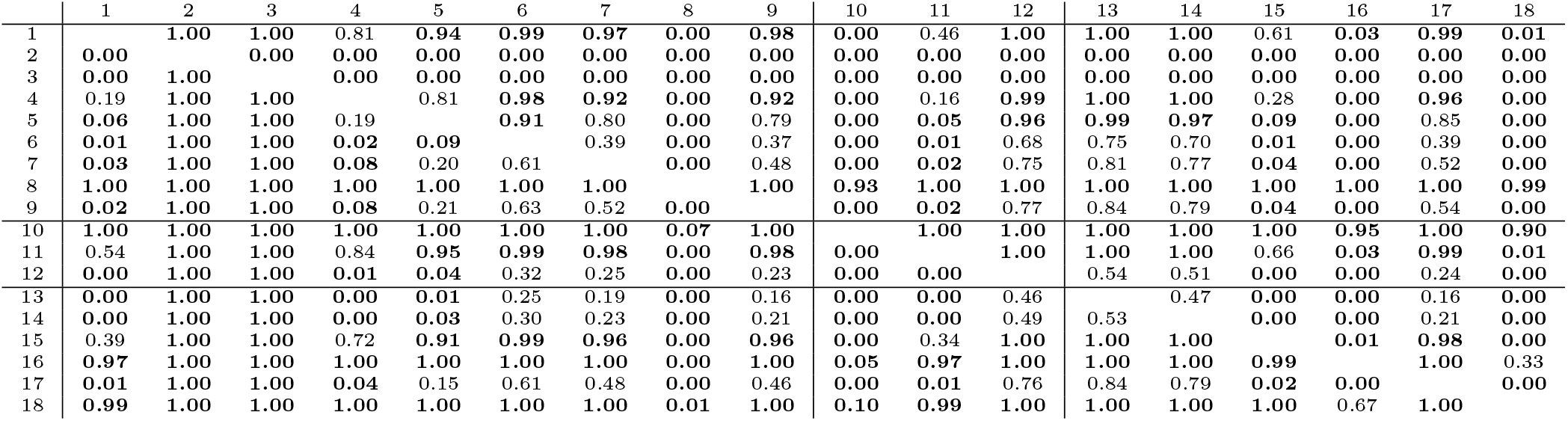
Pairwise contrasts 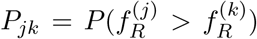 of maximum growth rates *f_R_*. Bold numbers and block partitioning as in Table 2.

The conversion factors *c_A_* and *c_R_* did not show significant pairwise differences for steady-state time series 1–9 (Fig. 4c,d; Tables 4,5, top left blocks; with few exceptions for *c_R_*). Some pairwise differences among cyclic time series 13–18 (bottom right blocks) and to steady-state time series (top right and bottom left blocks) were observed, without being as systematic as in *f_A_*. The uncertainty in parameter estimates is slightly larger in steady-state time series than in cycles time series, but the effect is not as prominent as in *f_A_* estimates (see also full tables of estimates, Tabs. A1–A4, Supporting Information). All findings of this section regarding uncertainties in the parameter estimation were in accordance to the simulation study’s results.

**Table 4:**
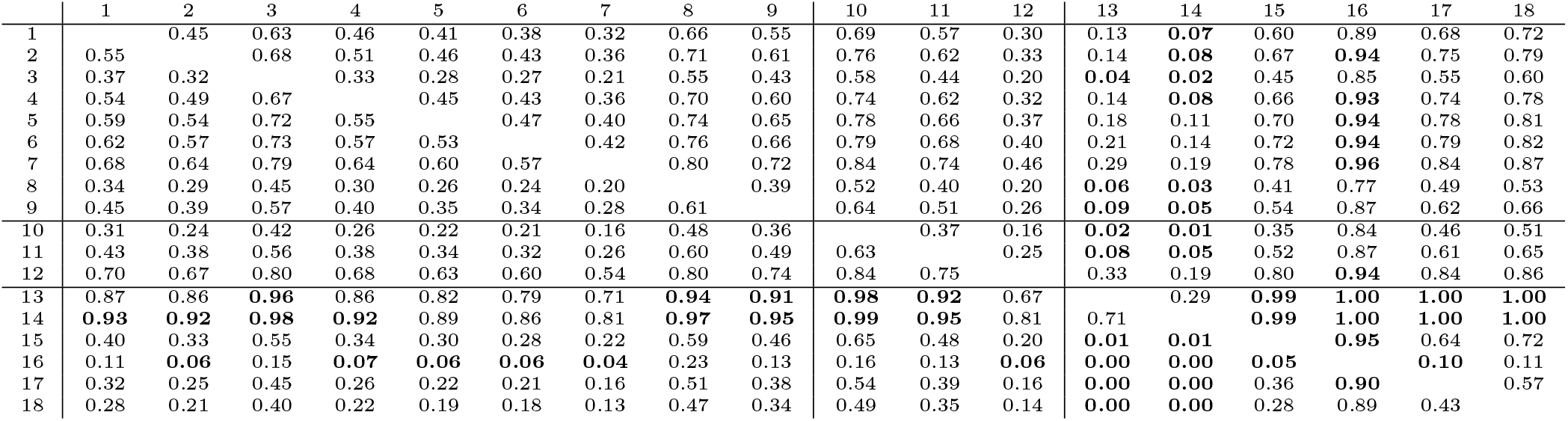
Pairwise contrasts 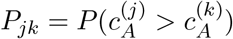 of conversion factors *c_A_*. Bold numbers and block partitioning as in Table 2.

**Table 5:**
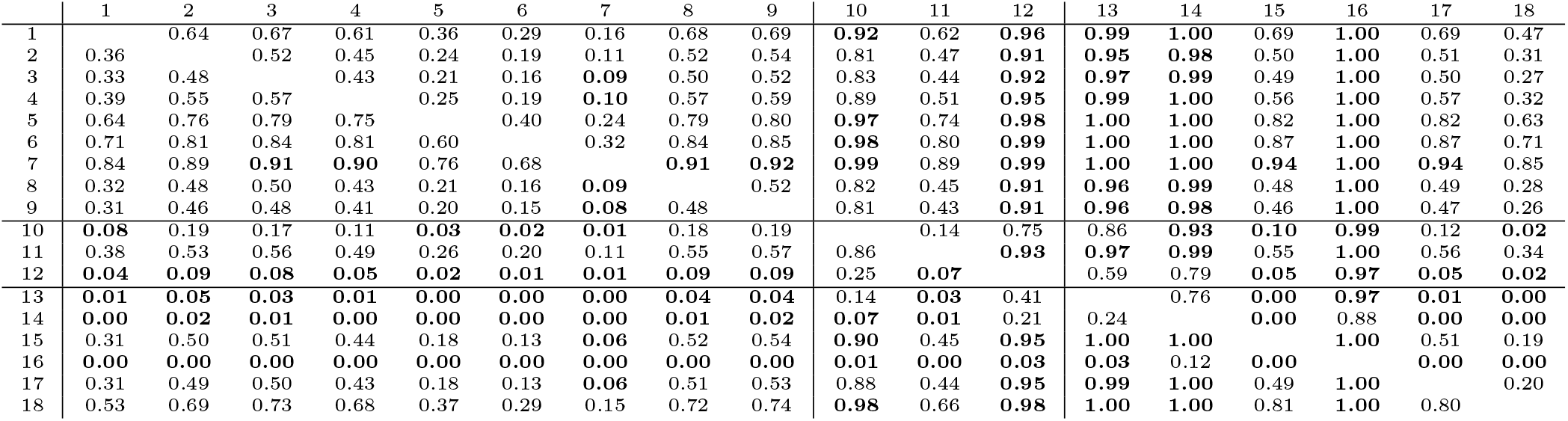
Pairwise contrasts 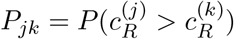 of conversion factors *c_R_*. Bold numbers and block partitioning as in Table 2.

### Comparison to complete pooling model fitting

To further support the importance of variation among the parameter estimates, we also fitted the complete pooling model using a single set of parameters {*f_A_, f_R_, c_A_, c_R_*} for all 18 time series as a null model. We used the same priors and model fitting specifications as above. Although a formal model comparison via information criteria is generally available for Bayesian statistics (Vehtari et al. 2018), it is not applicable to our dynamical model, since predictions (and therefore residuals) are correlated along time. Hence, we compare the complete pooling and the partial pooling (hierarchical) model qualitatively and quantitatively via their predictions. In contrast to the previous hierarchical model, the complete pooling model produces transient and steady-state predictions for all time series. Cyclic dynamics can not be reproduced (Supporting Information, Figs. A8&A9). Also, the predicted equilibrium fails to reproduce the correct level of algae and rotifer states in some cases. This corresponds to a generally lower predictive accuracy when fitting algal and rotifers time series with the complete pooling model as compared to the hierachical model (Supporting Information, Fig. A7). The effect is highest for cyclic time series 13–18, but also pronounced in time series 2, 3, 8 and 10, where the equilibrium states are not reproduced correctly.

### Transition from cyclic dynamics to steady states

To support the observation that low maximum growth rates *f_A_* cause cyclic dynamics while high maximum growth rates yield a steady-state equilibrium (Fig. 4), we performed a simulation study. We numerically simulated the chemostat model (eqns. 1–3) while systematically varying growth rates *f_A_* between 1 d^−1^ and 7.389 d^−1^ (corresponding to parameter values ln(*f_A_*) between 0 and 2). The maximum growth rate of the predator and the conversion factors were held constant at their estimated overall means exp(*μ*_ln(*θ*)_) across the 18 time series (*f_R_* = 1.419 d^−1^, *c_A_* = 3.865 · 10^7^ cells μmol^−1^, *c_R_* = 1.065 · 10^−5^ ind cells^−1^). The bifurcation diagram (Fig. 5) shows system states of simulations over 200 days after discarding the first 100 days and clearly indicates cycles for low growth rates and steady states for high growth rates, with the Hopf bifurcation located in *f_A_* = 4.446 d^−1^.

**Figure 5:**
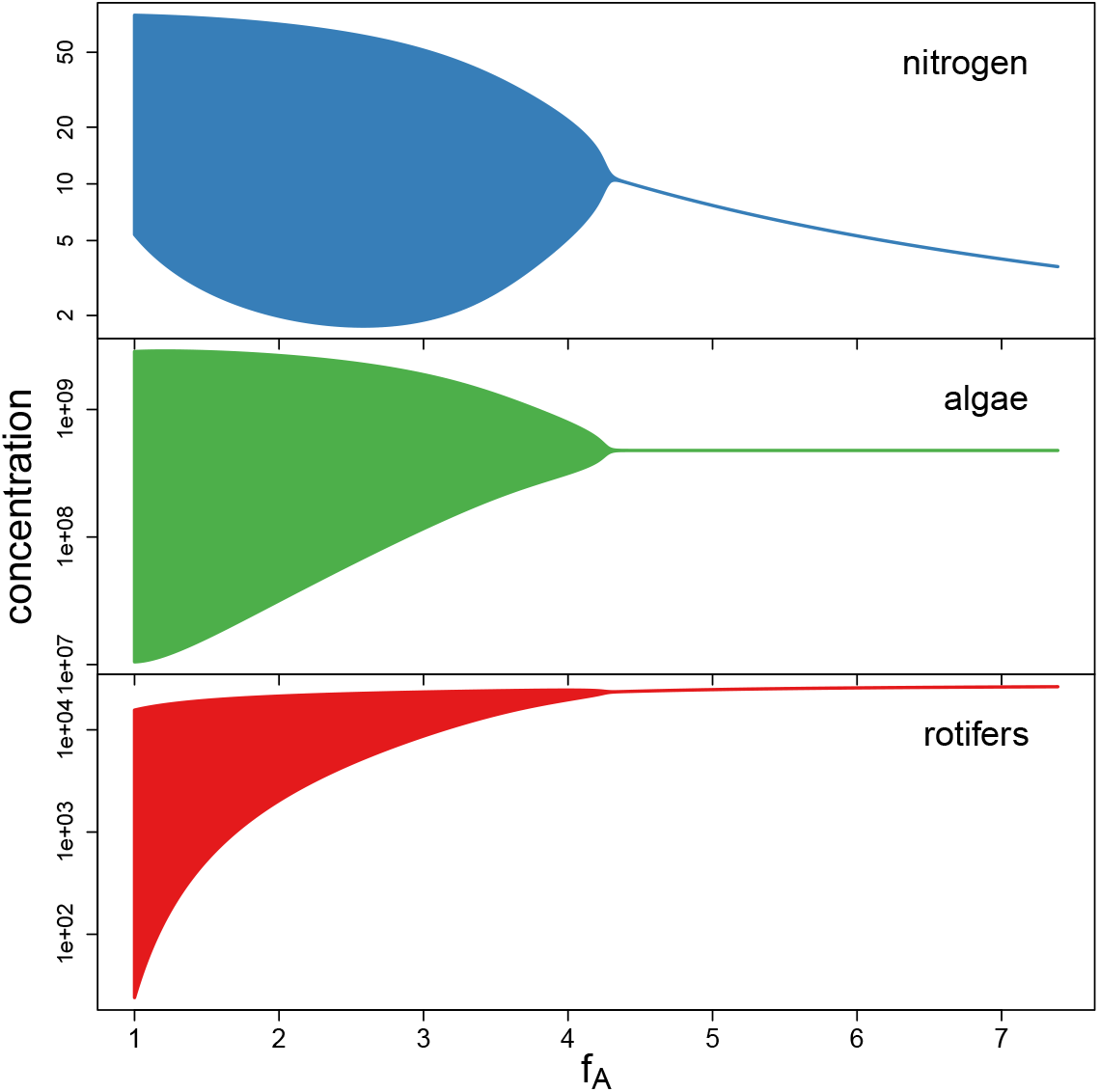
Bifurcation diagram for simulations with varying maximum growth rate *f_A_*, while keeping the remaining parameters constant at the fitted overall means. A Hopf bifurcation occurs at *f_A_* = 4.446.

## Discussion

In this study we presented how a differential equations model can be fitted to observed time series data of species abundances, taking predator-prey dynamics as an example. Next to obtaining key model parameters in situ, this method allows to decipher variability in the outcomes among replicates and to point towards probable sources of this variability.

The comparison of choosing the time series identity as a random effect (partial pooling) to a model using only a single set of parameters (complete pooling) shows that allowing for such a variability in the parameters between replicates can be crucial to see whether time series data agree with a model. Especially if a certain parameter is close to a bifurcation, as it seems to be the case in our system for the maximum growth rate of the algae *f_A_*, minor deviations in this parameter result in different predictions for the system dynamics. In such cases, in-situ parameter estimation allows the detection of parameter sets for both steady states and cyclic dynamics, which can be separated by a multi-dimensional bifurcation boundary in models with a high number of parameters. Thereby, the chosen model structure may be accepted for all replicates, even if their dynamics differ, without increasing the model complexity. Instead, more than one parameter set might be needed to cover the whole diversity of possible and observable patterns in population dynamics.

Of course, this method requires that the model structure used for fitting the data applies sufficiently well to the mechanisms acting in the system delivering the data. Also, the data quality has to be high enough. We see that the dynamics of the prey are fitted better than those of the predator, which may be explained by different mechanisms: (i) The densities of the prey in the data are much larger than those of the predator. Assuming that the relative experimental error of counting decreases with larger individual numbers, this error should be smaller for the prey than for the predator. This increases the regularity of the dynamical patterns and simplifies the fitting of the prey time series. (ii) Rotifers, as metazoan animals, possess a more complex life-cycle than algae. Their dynamics may be affected by age-structure, variable resource co-limitation and other factors, which, for simplicity, were not included in the model. These non-modelled processes obviously decrease the goodness of fit. (iii) As no data was available for the resource of the prey, the parameters of the prey were confined only by the prey data, leaving more flexibility to improve the fitting in the prey’s states. Contrarily, the parameters of the predator were more restricted, as both prey and predator data was available, leaving less flexibility for fitting the predator’s states.

Interestingly, the type of population dynamics affects the quality of the inference as well. Population cycles contain a higher degree of information on the system than steady states, as also rates of change are apparent, additional to the biomasses of the trophic levels. This manifests in more narrow parameter estimates (Fig. 4) and leads to more certain predictions (Figs. 2,3). Also, steady-state fits give rather (and partially unre-alistically) high estimates for algal growth rates, whereas those from cyclic population dynamics are estimated close to published values. These findings were also confirmed in a simulation study by fitting the model to synthetic data generated by known parameters (Fig. 1).

Variability in key parameters suggests a heterogeneity in the traits that are encoded by these parameters. Heterogeneous traits imply intra-specific variability which may enable populations to escape perturbations and to persist in the presence of strong stressors (Reusch et al. 2005; Bell and Gonzalez 2009; Chevin et al. 2010). Bayesian parameter inference provides uncertainty estimates on key parameters and thereby allows to detect such variability in experiments. We propose that this technique may also help to quantify the presence of trait heterogeneity in nature.

## Acknowledgements

We thank Bernd Blasius for his contribution to the acquisition and provision of the data.

## Authors’ contributions

BR, UG and MR conceived the ideas, GW and GF contributed the data, BR developed the methods, BR and MR led the writing of the manuscript. All authors contributed critically to the drafts and gave final approval for publication.

## Supplement

### Stan model code

~~~
functions{
   // ODE right hand side
   real [] frmodel(real t, real [] B, real [] p, real [] x_r, int [] x_i) {
     // p[1] S0, inflow concentration
     // p[2] delta, inflow rate
     // p [3] fA, max. growth rate
     // p[4] hA, half - saturation density
     // p[5] 1/cA, conversion factor
     // p[6] fR, max. growth rate
     // p[7] hR, half - saturation density
     // p [8] 1/cR, conversion factor
     real dBdt [3];
     real feeding_S;
     real feeding_A;
     if (B[1]<1e-6 || B[1]>1e16 || B[2]<1e-6 || B[2]>1e16 || B[3]<1e-6 || B[3]>1e16 ){
        dBdt [1] = 0;
        dBdt [2] =0;
        dBdt [3] =0;
     }
     else{
       feeding_S = (p[3]/(p[4]+B[1]))*B [1]*B[2];
       feeding_A = (p[6]/(p[7]+B[2]))*B [2]*B[3];
       dBdt [1] = (p[1]-B [1])*p [2] - p [5]*feeding_S;
       dBdt [2] = feeding_S - p [8]*feeding_A - p[2]*B[2];
       dBdt [3] = feeding_A                         - p[2]*B[3];
     }
     return dBdt;
   }
}
data{
     int m;                     // number of time series
     int n[m];                  // lengths of the m time series
     int nmax;                  // max(n)
     real time0[m];             // initial timepoints of the m time series
     real logA0[m];             // initial algal densities (log)
     real logR0[m];             // initial rotifer densities (log)
     real time [m,nmax];        // each row i contains times of observations and is filled with placeholder 0s for n[i]<j<=nmax
     real logA [m,nmax];        // algal observations, placeholders as in time[,]
     real logR [m,nmax];        // rotifer observations, placeholders as in time[,]
}
transformed data {       // just empty placeholders for function call
    real x_r [0];
    int x_i [0];
}
parameters {
     real mu_fA;
     real<lower=0> sigma_fA;
     real fA[m]; // maximum growth rates, A <-S real mu_fR;
     real<lower=0> sigma_fR;
     real fR[m]; // maximum growth rates, R <-A real mu_cA;
     real<lower=0> sigma_cA;
     real cA[m]; // conversion factors, A <-S real mu_cR;
     real<lower=0> sigma_cR;
     real cR[m]; // conversion factors, R <-A
     real<lower=0> sdev_A; // standard deviation of residuals
     real<lower=0> sdev_R; // standard deviation of residuals
     real logB0[m,3]; // initial values for ODE
}
model {
     // intermediate parameters
     real p[8]; // hand over to ODE
     real Bsim [nmax,3]; // simulated values, matrix. dim1 = time, dim2 = dim_ODE = 3 real B0[3]; // initial values for ODE
     // priors (informative) mu_fA ~ normal(1.591, 1.0); sigma_fA ~ normal(0, 1.0);
     mu_fR ~ normal(0.356, 1.0); sigma_fR ~ normal(0, 1.0);
     mu_cA ~ normal(17.190, 1.0); sigma_cA ~ normal(0, 1.0);
     mu_cR ~ normal(−11.067, 1.0); sigma_cR ~ normal(0, 1.0);
     for(j in 1:m){
       fA[j] ~ normal (mu_fA, sigma_fA);
       fR[j] ~ normal(mu_fR, sigma_fR);
       cA[j] ~ normal(mu_cA, sigma_cA);
       cR[j] ~ normal(mu_cR, sigma_cR);
       logB0 [j, 1] ~ normal(2,2);
       logB0 [j, 2] ~ normal(20,2);
       logB0 [j, 3] ~ normal(10,2);
    }
    sdev_A ~ normal(0, 2);
    sdev_R ~ normal(0, 2);
    // likelihood
    for (j in 1: m){             // loop over time series
       p[1] = 80.0;              // S0, inflow concentration
       p[2] = 0.55;              // delta, inflow rate
       p[3] = exp(fA[j]);        // max. growth rate
       p[4] = 4.3;               // half - saturation density
       p[5] = 1/exp(cA[j]);      // conversion factor
       p[6] = exp(fR[j]);        // max. growth rate
       p[7] = 7.5e8;             // half - saturation density
       p[8] = 1/exp(cR[j]);      // conversion factor
       // ODE simulation
       for (i in 1: 3) {
         B0[i] = exp(logB0 [j,i]);
       }
       Bsim[1:n[j], 1:3] = integrate_ode_rk45(frmodel,B0,time0 [j],time [j,1:n[j]],p,x_r,x_i);
       // residuals
       logA0[j] ~ normal(logB0[j,2], sdev_A);
       logR0[j] ~ normal(logB0[j,3], sdev_R);
       for (i in 1:n[j]){
         logA[j,i] ~ normal(log(Bsim[i,2]), sdev_A);
         logR[j,i] ~ normal(log(Bsim[i,3]), sdev_R);
       }
    } // end for (j in 1: m)
}
~~~

**Figure A1:**
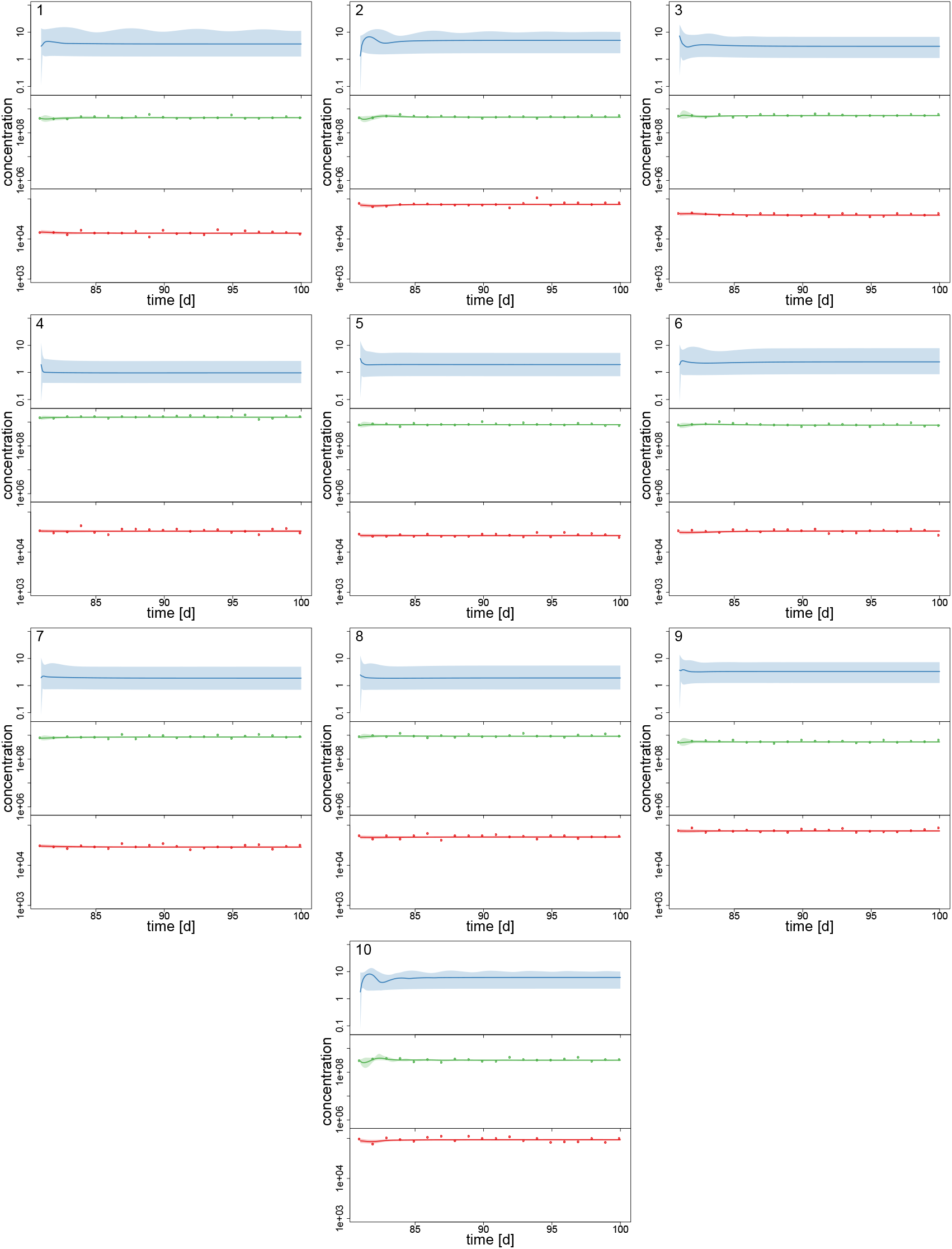
**Simulated time series** 1–10 of 20, data (dots) and posterior predictions of fitted hierarchical model for nitrogen (blue), algae (green) and rotifers (red). Solid lines represent median predictions, shaded areas depict 95% highest density intervals of the predictions.

**Figure A2:**
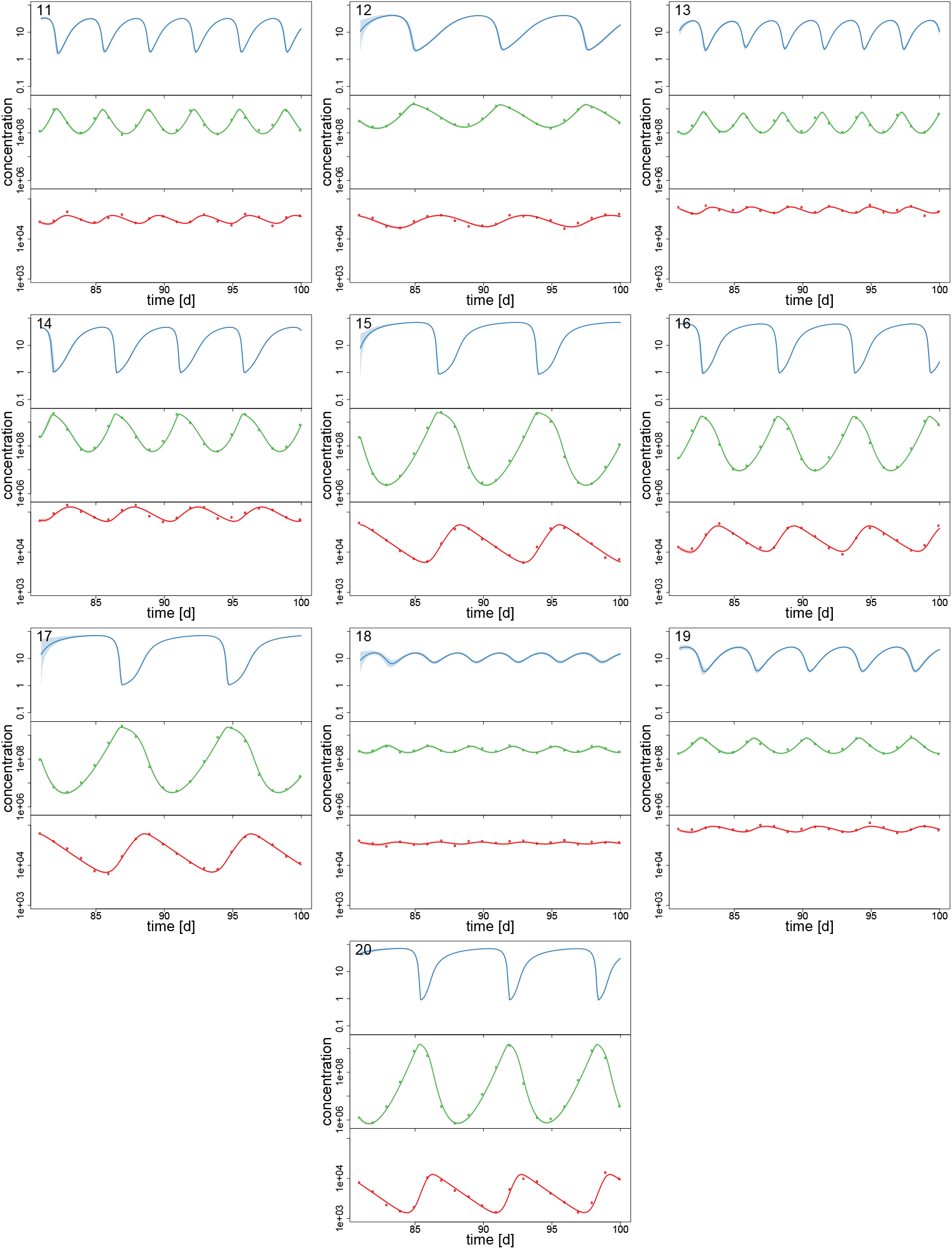
**Simulated time series** 11–20 of 20, data (dots) and posterior predictions of fitted hierarchical model for nitrogen (blue), algae (green) and rotifers (red). Solid lines represent median predictions, shaded areas depict 95% highest density intervals of the predictions.

**Figure A3:**
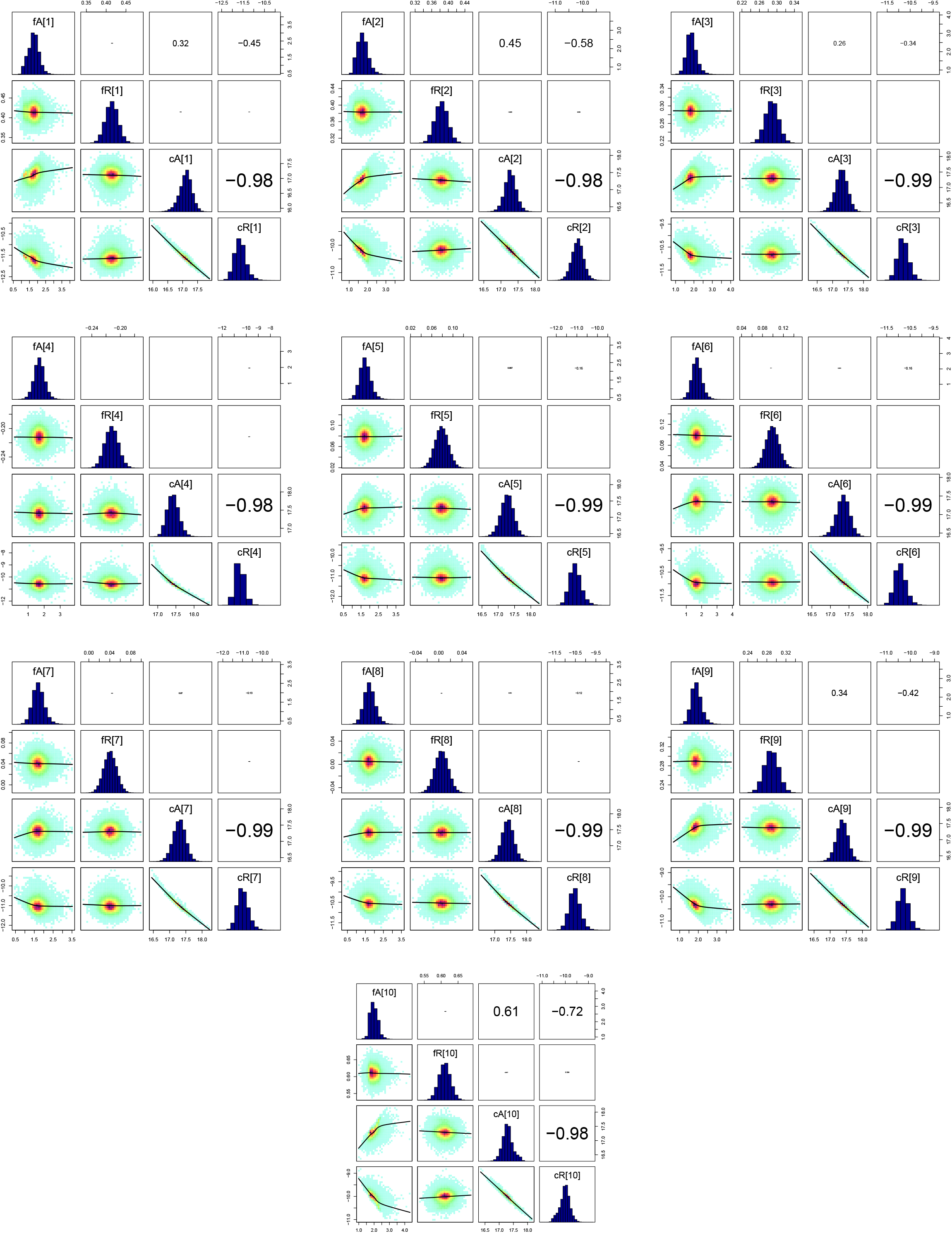
Pairs plots of **simulated time series** 1–10 of 20 parameters (on ln-scale). Marginal posterior densities (diagonal panels), pairwise posterior densities (lower left panels, red indicates (high density, black line represents nonlinear fit of the correlation) and correlation coefficients (upper right panels, fontsize indicates high correlation).

**Figure A4:**
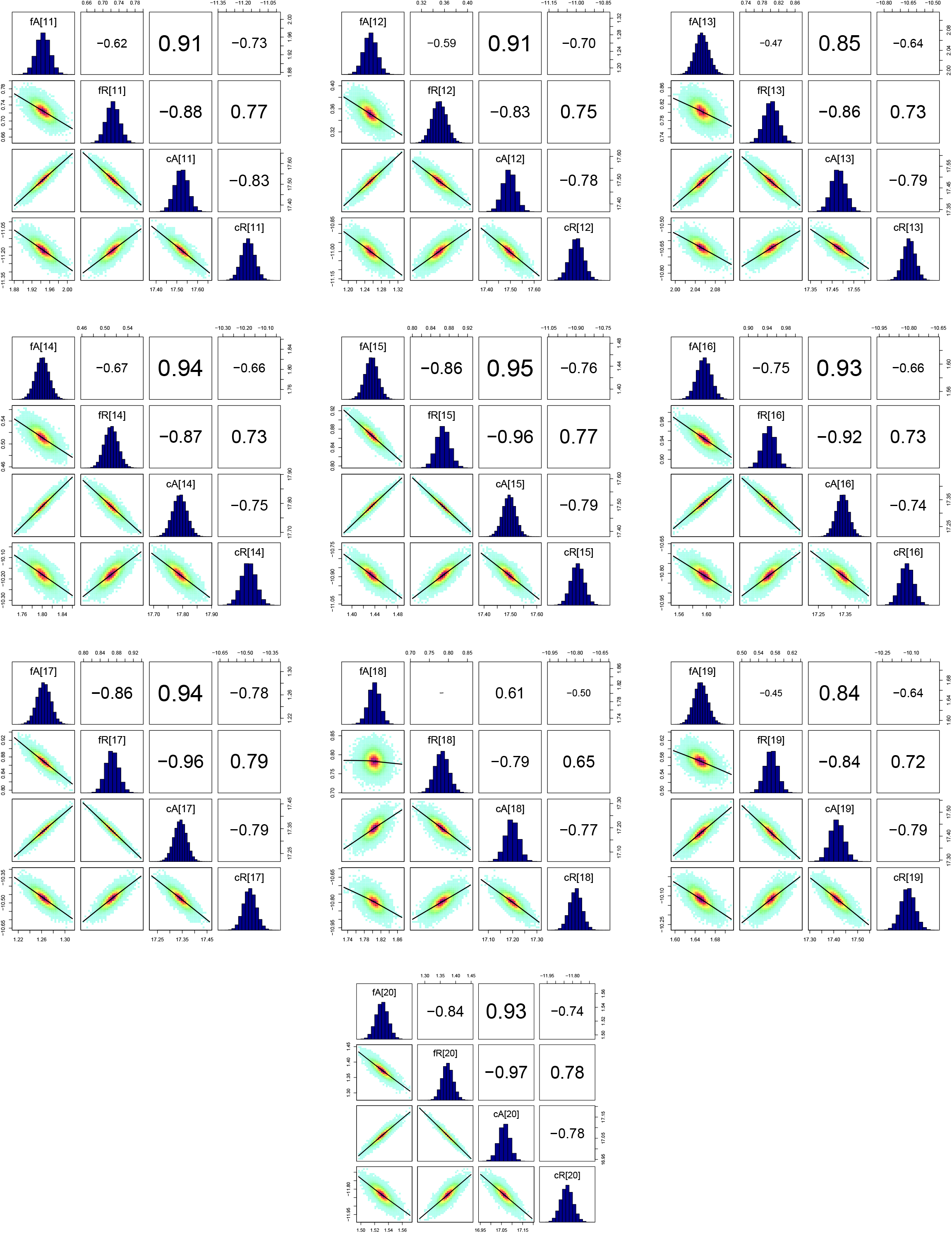
Pairs plots of **simulated time series** 11–20 of 20 parameters (on ln-scale). Panels as in Fig. A3.

**Table A1:**
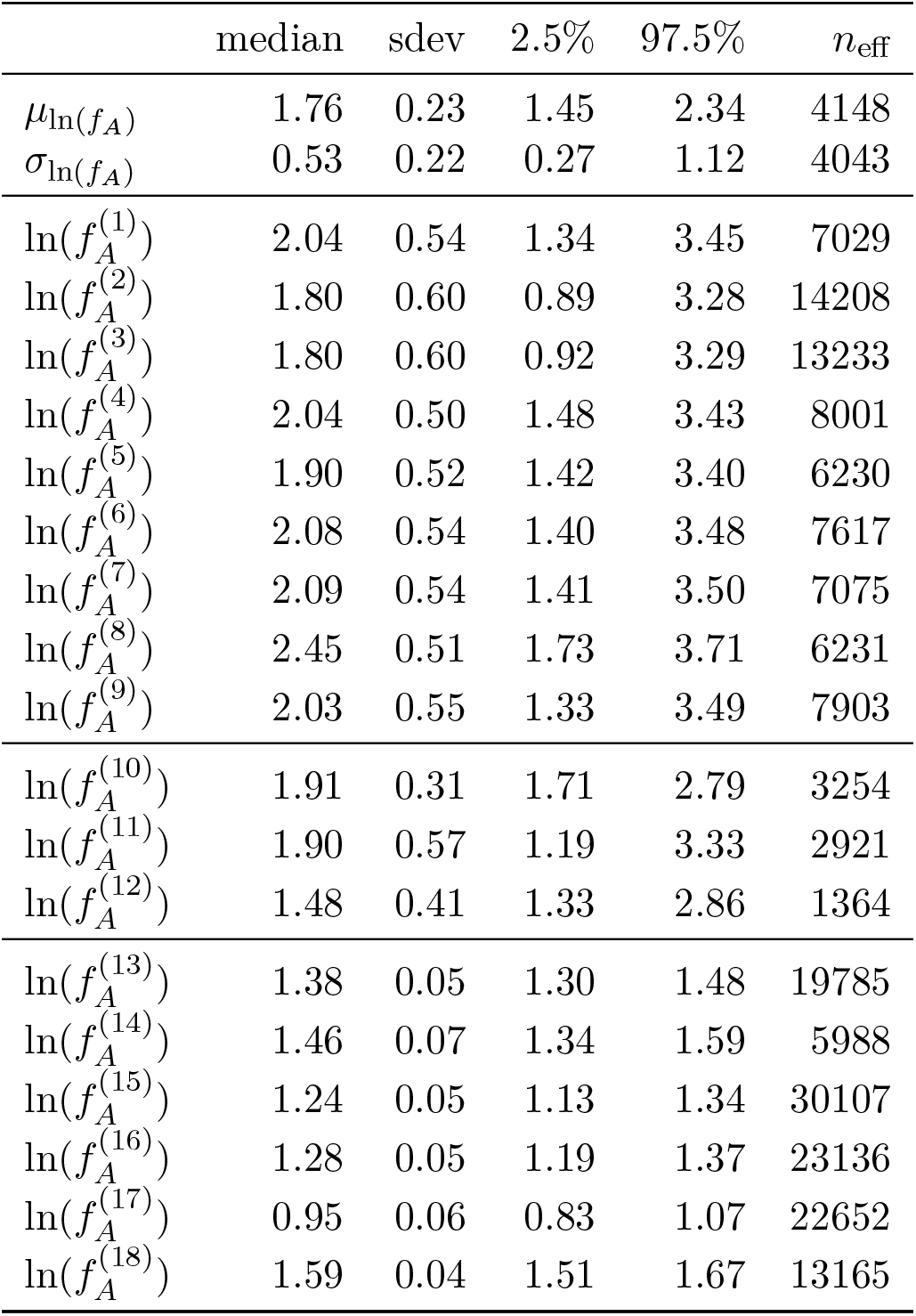
Summary statistics for maximum growth rate *f_A_* [d^−1^] in hierarchical model for experimental chemostat data. Gelman-Rubin statistics *Ȓ* < 1.01 for all parameters verify convergence, *n*_eff_ is the number of independent samples. Block partitioning refers to time series 1–9 featuring a steady-state equilibrium, 10–12 having multimodal posterior distributions featuring both steady states and cycles, and 13–18 featuring cyclic dynamics.

**Table A2:** Summary statistics for maximum growth rate *f_R_*[d^−1^] in hierarchical model for experimental chemostat data. Gelman-Rubin statistics *Ȓ* < 1.01 for all parameters verify convergence, *n_eff_* is the number of independent samples. Block partitioning as in Table A1.

| | median | sdev | 2.5% | 97.5% | $n_{\text{eff}}$ |
| --- | --- | --- | --- | --- | --- |
| $\mu_{\ln(f_R)}$ | 0.35 | 0.06 | 0.23 | 0.46 | 47132 |
| $\sigma_{\ln(f_R)}$ | 0.23 | 0.05 | 0.16 | 0.34 | 41793 |
| $\ln(f_R^{(1)})$ | 0.42 | 0.05 | 0.32 | 0.52 | 52007 |
| $\ln(f_R^{(2)})$ | -0.13 | 0.03 | -0.20 | -0.06 | 49242 |
| $\ln(f_R^{(3)})$ | 0.06 | 0.03 | 0.01 | 0.12 | 54068 |
| $\ln(f_R^{(4)})$ | 0.37 | 0.03 | 0.30 | 0.43 | 48709 |
| $\ln(f_R^{(5)})$ | 0.33 | 0.03 | 0.27 | 0.39 | 62015 |
| $\ln(f_R^{(6)})$ | 0.26 | 0.04 | 0.19 | 0.34 | 55771 |
| $\ln(f_R^{(7)})$ | 0.28 | 0.05 | 0.19 | 0.38 | 58240 |
| $\ln(f_R^{(8)})$ | 0.76 | 0.06 | 0.64 | 0.88 | 54787 |
| $\ln(f_R^{(9)})$ | 0.28 | 0.05 | 0.19 | 0.38 | 54633 |
| $\ln(f_R^{(10)})$ | 0.66 | 0.04 | 0.58 | 0.74 | 39719 |
| $\ln(f_R^{(11)})$ | 0.43 | 0.05 | 0.33 | 0.53 | 50021 |
| $\ln(f_R^{(12)})$ | 0.23 | 0.05 | 0.13 | 0.32 | 2738 |
| $\ln(f_R^{(13)})$ | 0.23 | 0.03 | 0.17 | 0.29 | 19431 |
| $\ln(f_R^{(14)})$ | 0.23 | 0.04 | 0.16 | 0.31 | 19802 |
| $\ln(f_R^{(15)})$ | 0.40 | 0.04 | 0.31 | 0.49 | 25485 |
| $\ln(f_R^{(16)})$ | 0.55 | 0.05 | 0.46 | 0.65 | 20477 |
| $\ln(f_R^{(17)})$ | 0.28 | 0.04 | 0.20 | 0.36 | 23168 |
| $\ln(f_R^{(18)})$ | 0.58 | 0.05 | 0.49 | 0.67 | 24308 |

**Table A3:** Summary statistics for conversion factor *c_A_*[cells μmol^−1^] in hierarchical model for experimental chemostat data. Gelman-Rubin statistics *Ȓ* < 1.01 for all parameters verify convergence, *n*_eff_ is the number of independent samples. Block partitioning as in Table A1.

| | median | sdev | 2.5% | 97.5% | $n_{\text{eff}}$ |
| --- | --- | --- | --- | --- | --- |
| $\mu_{\ln(c_A)}$ | 17.47 | 0.08 | 17.30 | 17.62 | 6021 |
| $\sigma_{\ln(c_A)}$ | 0.19 | 0.06 | 0.11 | 0.33 | 12142 |
| $\ln(c_A^{(1)})$ | 17.46 | 0.19 | 17.08 | 17.81 | 13089 |
| $\ln(c_A^{(2)})$ | 17.49 | 0.16 | 17.17 | 17.82 | 17147 |
| $\ln(c_A^{(3)})$ | 17.39 | 0.15 | 17.08 | 17.68 | 16103 |
| $\ln(c_A^{(4)})$ | 17.49 | 0.17 | 17.13 | 17.78 | 13885 |
| $\ln(c_A^{(5)})$ | 17.51 | 0.17 | 17.15 | 17.84 | 11633 |
| $\ln(c_A^{(6)})$ | 17.52 | 0.18 | 17.16 | 17.90 | 12093 |
| $\ln(c_A^{(7)})$ | 17.56 | 0.19 | 17.20 | 17.95 | 11531 |
| $\ln(c_A^{(8)})$ | 17.37 | 0.20 | 16.93 | 17.73 | 9904 |
| $\ln(c_A^{(9)})$ | 17.43 | 0.18 | 17.05 | 17.78 | 14406 |
| $\ln(c_A^{(10)})$ | 17.36 | 0.13 | 17.09 | 17.63 | 10915 |
| $\ln(c_A^{(11)})$ | 17.42 | 0.17 | 17.08 | 17.76 | 8854 |
| $\ln(c_A^{(12)})$ | 17.62 | 0.19 | 17.13 | 17.84 | 1198 |
| $\ln(c_A^{(13)})$ | 17.68 | 0.08 | 17.52 | 17.84 | 16372 |
| $\ln(c_A^{(14)})$ | 17.74 | 0.10 | 17.54 | 17.94 | 8589 |
| $\ln(c_A^{(15)})$ | 17.41 | 0.08 | 17.25 | 17.58 | 22047 |
| $\ln(c_A^{(16)})$ | 17.21 | 0.10 | 17.02 | 17.39 | 17519 |
| $\ln(c_A^{(17)})$ | 17.37 | 0.09 | 17.20 | 17.54 | 17976 |
| $\ln(c_A^{(18)})$ | 17.35 | 0.07 | 17.21 | 17.50 | 15892 |

**Table A4:** Summary statistics for conversion factor *c_R_*[ind cells^−1^] in hierarchical model for experimental chemostat data. Gelman-Rubin statistics *Ȓ* < 1.01 for all parameters verify convergence, *n*_eff_ is the number of independent samples. Block partitioning as in Table A1.

| | median | sdev | 2.5% | 97.5% | $n_{\text{eff}}$ |
| --- | --- | --- | --- | --- | --- |
| $\mu_{\ln(c_R)}$ | -11.44 | 0.12 | -11.67 | -11.17 | 8052 |
| $\sigma_{\ln(c_R)}$ | 0.37 | 0.09 | 0.23 | 0.59 | 12030 |
| $\ln(c_R^{(1)})$ | -11.26 | 0.24 | -11.70 | -10.77 | 10999 |
| $\ln(c_R^{(2)})$ | -11.38 | 0.27 | -11.87 | -10.78 | 16003 |
| $\ln(c_R^{(3)})$ | -11.39 | 0.23 | -11.79 | -10.91 | 14939 |
| $\ln(c_R^{(4)})$ | -11.35 | 0.20 | -11.68 | -10.89 | 12647 |
| $\ln(c_R^{(5)})$ | -11.17 | 0.21 | -11.56 | -10.72 | 10548 |
| $\ln(c_R^{(6)})$ | -11.10 | 0.23 | -11.55 | -10.63 | 10967 |
| $\ln(c_R^{(7)})$ | -10.97 | 0.24 | -11.44 | -10.48 | 10475 |
| $\ln(c_R^{(8)})$ | -11.39 | 0.24 | -11.82 | -10.85 | 8885 |
| $\ln(c_R^{(9)})$ | -11.40 | 0.24 | -11.83 | -10.89 | 13009 |
| $\ln(c_R^{(10)})$ | -11.61 | 0.16 | -11.96 | -11.31 | 8269 |
| $\ln(c_R^{(11)})$ | -11.35 | 0.24 | -11.79 | -10.85 | 3749 |
| $\ln(c_R^{(12)})$ | -11.79 | 0.19 | -12.07 | -11.30 | 2700 |
| $\ln(c_R^{(13)})$ | -11.82 | 0.11 | -12.04 | -11.61 | 20815 |
| $\ln(c_R^{(14)})$ | -11.94 | 0.14 | -12.20 | -11.67 | 14453 |
| $\ln(c_R^{(15)})$ | -11.38 | 0.12 | -11.62 | -11.13 | 26505 |
| $\ln(c_R^{(16)})$ | -12.15 | 0.13 | -12.41 | -11.89 | 23006 |
| $\ln(c_R^{(17)})$ | -11.38 | 0.14 | -11.65 | -11.11 | 21300 |
| $\ln(c_R^{(18)})$ | -11.24 | 0.11 | -11.46 | -11.02 | 19116 |

**Figure A5:**
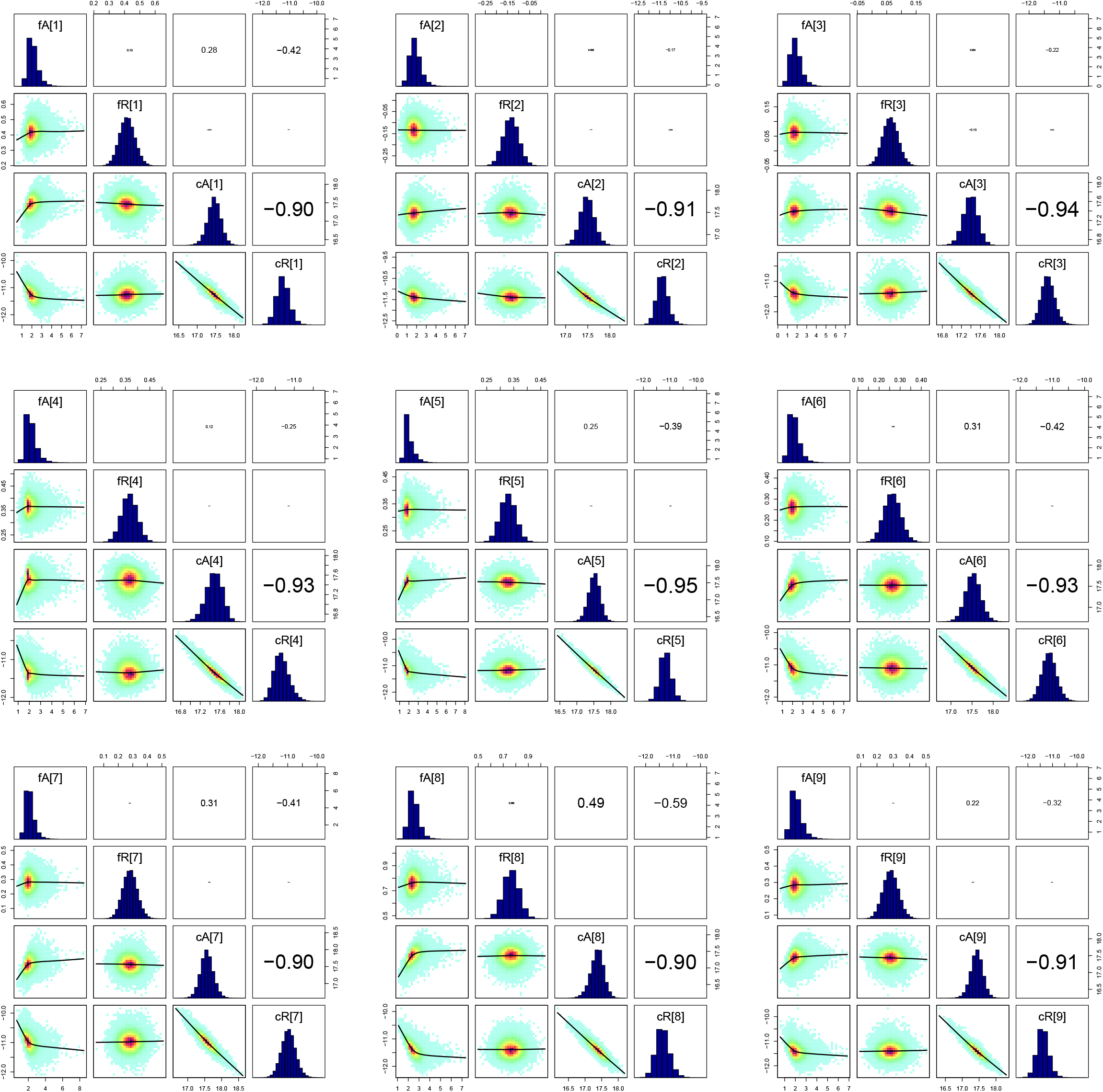
Pairs plots of **experimental time series** 1–9 of 18 parameters (on ln-scale). Panels as in Fig. A3.

**Figure A6:**
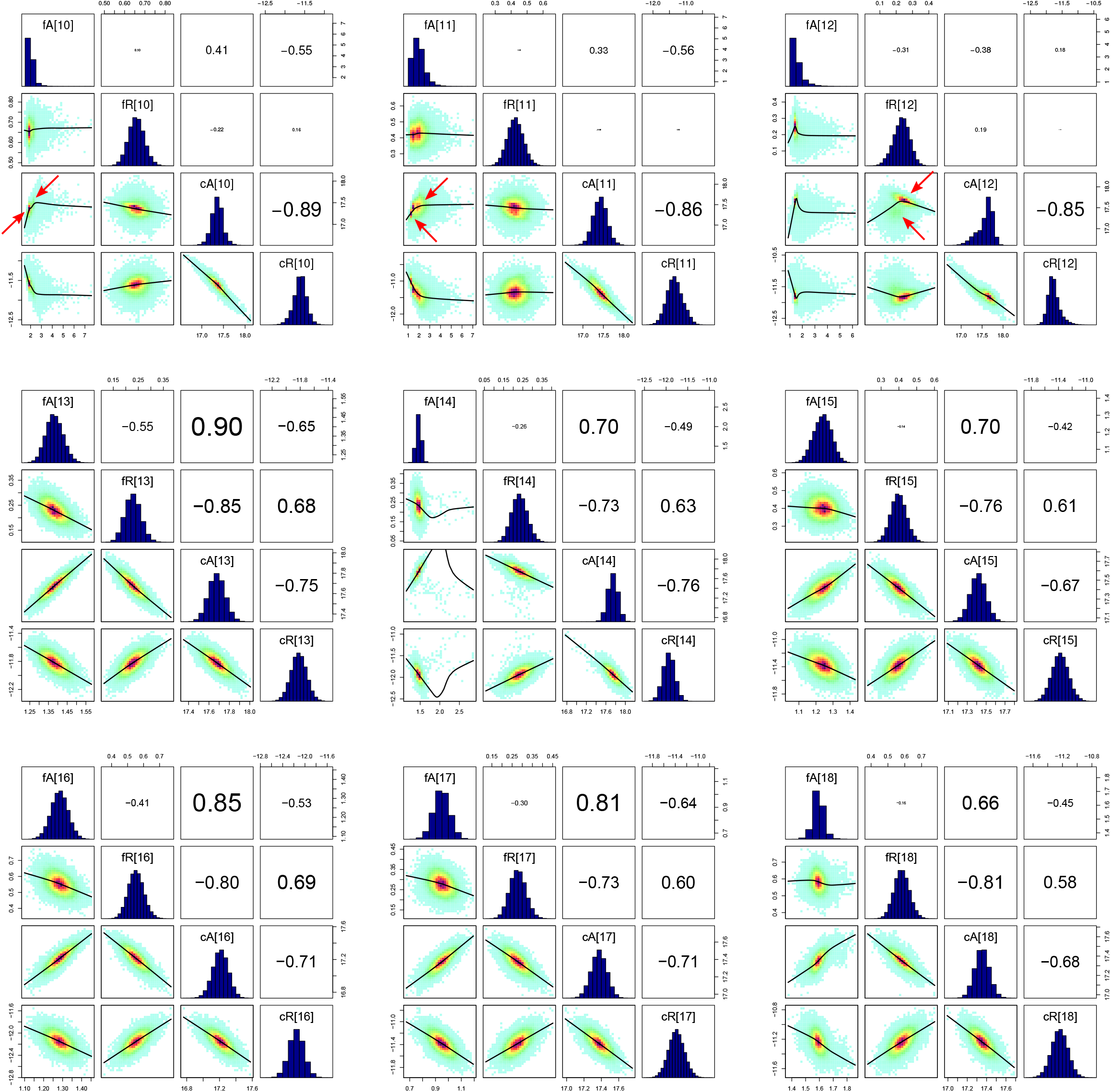
Pairs plots of **experimental time series** 10–18 of 18 parameters (on ln-scale). Panels as in Fig. A3. Red arrows indicate multimodalities in time series 10–12.

**Figure A7:**
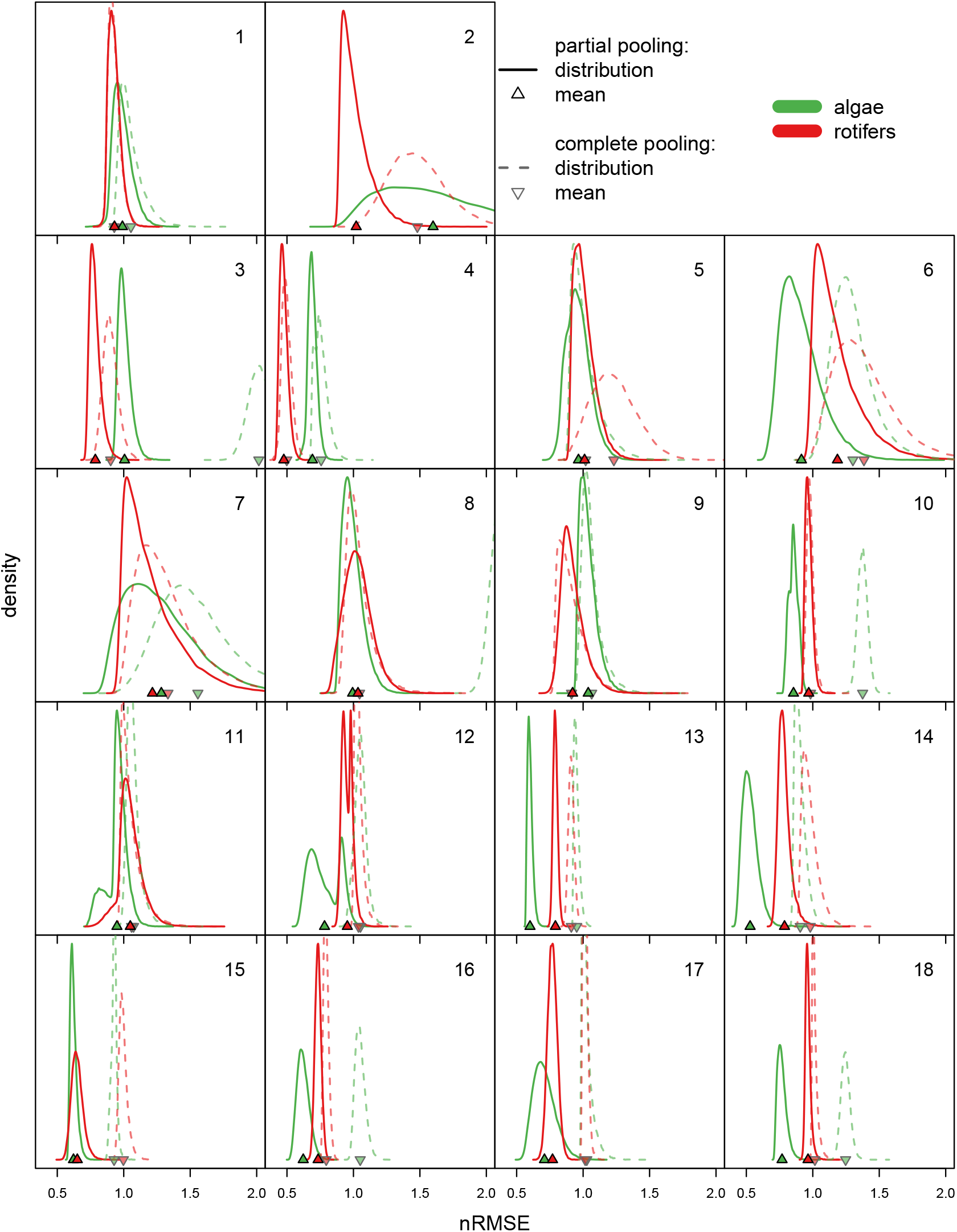
Predictive accuracy assessed by posterior distributions of the normalized root-mean-square error (nRMSE) 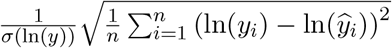 with log-scaled data ln(*y_i_*) and predictions ln(*ŷ_i_*), normalized by each time series’ empirical standard deviation *σ*(ln(*y*)), *y* = *A*^(*j*)^, *R*^(*j*)^, *j* = 1,…, 18. Algae (green) and rotifer (red) nRMSE for partial pooling model using distinct parameters for each time series (solid lines) and complete pooling model using the same parameter set across all time series (dashed lines).

**Table A5:** Summary statistics for complete pooling model for experimental chemostat data. Gelman-Rubin statistics *Ȓ* < 1.01 for all parameters verify convergence, *n*_eff_ is the number of independent samples.

| | median | sdev | 2.5% | 97.5% | $n_{\text{eff}}$ |
| --- | --- | --- | --- | --- | --- |
| $\ln(f_A)$ | 1.40 | 0.80 | 0.22 | 3.23 | 36982 |
| $\ln(f_R)$ | 0.34 | 0.02 | 0.31 | 0.37 | 47063 |
| $\ln(c_A)$ | 15.75 | 0.04 | 15.67 | 15.83 | 29223 |
| $\ln(c_R)$ | -7.53 | 0.34 | -8.12 | -6.80 | 29730 |

**Figure A8:**
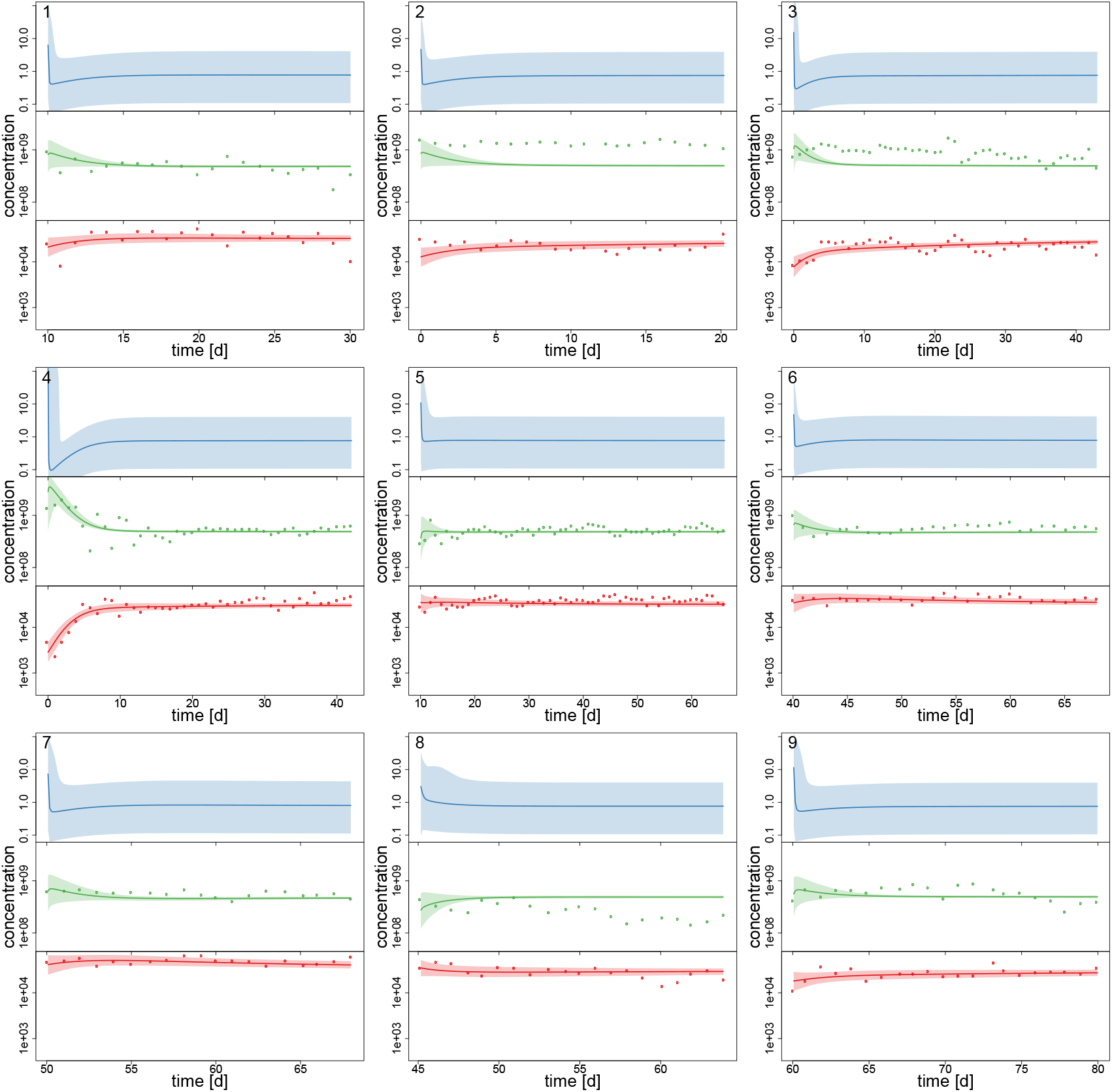
Experimental time series 1–9 of 18, data (dots) and posterior predictions of complete pooling model for nitrogen [μmol l^−1^] (blue), algae [cells l^−1^] (green) and rotifers [ind l^−1^] (red). Solid lines represent median predictions, shaded areas depict 95% highest density intervals of the predictions.

**Figure A9:**
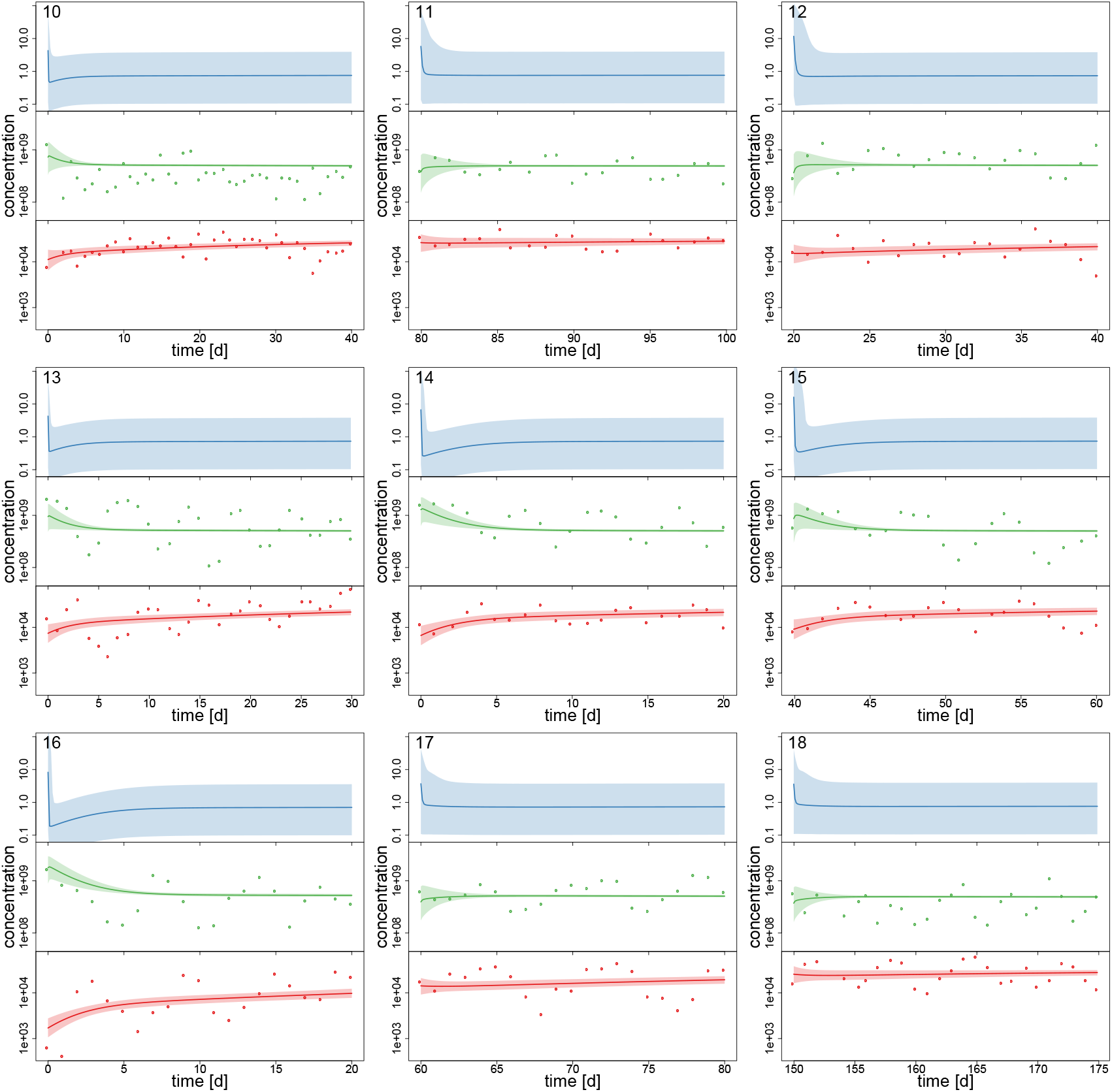
Experimental time series 10–18 of 18, data (dots) and posterior predictions of complete pooling model for nitrogen [μmol l^−1^] (blue), algae [cells l^−1^] (green) and rotifers [ind l^−1^] (red). Solid lines represent median predictions, shaded areas depict 95% highest density intervals of the predictions.

## References

Abrams, P. A. 1999. Is predator-mediated coexistence possible in unstable systems? Ecology 80: 608–621.

Almaraz, P. and Oro, D. 2011. Size-mediated non-trophic interactions and stochastic predation drive assembly and dynamics in a seabird community. Ecology 92: 1948–1958.

Almaraz, P. et al. 2012. Estimating partial observability and nonlinear climate effects on stochastic community dynamics of migratory waterfowl. J. Anim. Ecol. 81: 1113–1125.

Aster, R. C. et al. 2012. Parameter estimation and inverse problems. 2nd edition. Academic Press.

Barraquand, F. et al. 2017. Moving forward in circles: challenges and opportunities in modelling population cycles. Ecol. Lett. 20: 1074–1092.

Becks, L. and Arndt, H. 2013. Different types of synchrony in chaotic and cyclic communities. Nat. Commun. 4: 1359.

Becks, L. et al. 2010. Reduction of adaptive genetic diversity radically alters eco-evolutionary community dynamics. Ecol. Lett. 13: 989–997.

Becks, L. et al. 2012. The functional genomics of an eco-evolutionary feedback loop: Linking gene expression, trait evolution, and community dynamics. Ecol. Lett. 15: 492–501.

Bell, G. and Gonzalez, A. 2009. Evolutionary rescue can prevent extinction following environmental change. Ecol. Lett. 12: 942–948.

Boersch-Supan, P. H. et al. 2017. deBInfer: Bayesian inference for dynamical models of biological systems in R. Methods Ecol. Evol. 8: 511–518.

Boit, A. and Gaedke, U. 2014. Benchmarking successional progress in a quantitative food eb. PLOS ONE 9: 1–25.

Boit, A. et al. 2012. Mechanistic theory and modelling of complex food-web dynamics in Lake Constance. Ecol. Lett. 15: 594–602.

Bolius, S. et al. 2017. High local trait variability in a globally invasive cyanobacterium. Freshwater Biology 62: 1879–1890.

Bolker, B. M. 2008. Ecological Models and Data in R. Princeton University Press.

Bolnick, D. I. et al. 2011. Why intraspecific trait variation matters in community ecology. Trends Ecol. Evol. 26: 183–192.

Cao, J. et al. 2008. Estimating a predator-prey dynamical model with the parameter cascades method. Biometrics 64: 959–967.

Carpenter, B. 2018. Predator-prey population dynamics: the Lotka-Volterra model in Stan. accessed August 13, 2018. eprint: http://mc-stan.org/users/documentation/case-studies/lotka-volterra-predator-prey.html.

Carpenter, B. et al. 2017. Stan: a probabilistic programming language. Journal of Statistical Software 76: 1–32.

Chevin, L.-M. et al. 2010. Adaptation, plasticity, and extinction in a changing environment: towards a predictive theory. PLOS Biology 8: 1–8.

Compagnoni, A. et al. 2016. The effect of demographic correlations on the stochastic population dynamics of perennial plants. Ecol. Monogr. 86: 480–494.

Cortez, M. H. 2018. Genetic variation determines which feedbacks drive and alter predator-prey eco-evolutionary cycles. Ecological Monographs 88: 353–371.

Costantino, R. F. et al. 2005. “Nonlinear stochastic population dynamics: the flour beetle tribolium as an effective tool of discovery”. In: Population Dynamics and Laboratory Ecology. Vol. 37. Advances in Ecological Research. Academic Press, pp. 101–141.

Curtsdotter, A. et al. 2018. Ecosystem function in predator-prey food webs-confronting dynamic models with empirical data. Journal of Animal Ecology: 1–15.

DeLong, J. P. et al. 2014. Predator–prey dynamics and the plasticity of predator body size. Functional ecology 28: 487–493.

Ehrlich, E. et al. 2017. Trait–fitness relationships determine how trade-off shapes affect species coexistence. Ecology 98: 3188–3198.

Elderd, B. D. and Miller, T. E. X. 2015. Quantifying demographic uncertainty: Bayesian methods for Integral Projection Models (IPMs). Ecological Monographs 86: 15–1526.1.

Fussmann, G. F. et al. 2000. Crossing the Hopf bifurcation in a live predator-prey system. Science 290: 1358–1360.

Fussmann, K. E. et al. 2017. Interactive effects of shifting body size and feeding adaptation drive interaction strengths of protist predators under warming. bioRxiv: eprint: https://www.biorxiv.org/content/early/2017/01/20/101675.full.pdf.

Gaedke, U. and Klauschies, T. 2017. Analyzing the shape of observed trait distributions enables a data-based moment closure of aggregate models. Limnol. Oceanogr. Methods 15: 979–994.

Gelman, A. and Hill, J. 2007. Data Analysis Using Regression and Multilevel/Hierarchical Models. Cambridge University Press.

Gilioli, G. et al. 2008. Bayesian inference for functional response in a stochastic predator-prey system. Bull. Math. Biol. 70: 358–381.

Hefley, T. J. et al. 2013. Fitting population growth models in the presence of measurement and detection error. Ecol. Modell. 263: 244–250.

Hillebrand, H. and Matthiessen, B. 2009. Biodiversity in a complex world: Consolidation and progress in functional biodiversity research. Ecol. Lett. 12: 1405–1419.

Hosack, G. R. et al. 2012. Estimating density dependence and latent population trajectories with unknown observation error: Estimating unknown observation error. Methods Ecol. Evol. 3: 1028–1038.

Iijima, H. et al. 2013. Estimation of deer population dynamics using a bayesian state-space model with multiple abundance indices. Jour. Wild. Mgmt. 77: 1038–1047.

Johnson, L. R. et al. 2013. Bayesian inference for bioenergetic models. Ecology 94: 882–894.

Kath, N. J. et al. 2018. Accounting for activity respiration results in realistic trophic transfer efficiencies in allometric trophic network (ATN) models. Theor. Ecol. 1–11.

Kindsvater, H. K. et al. 2018. Overcoming the data crisis in biodiversity conservation. Trends Ecol. Evol.

Koons, D. N. et al. 2015. Disentangling the effects of climate, density dependence, and harvest on an iconic large herbivore’s population dynamics. Ecol. Appl. 25: 956–967.

Litchman, E. and Klausmeier, C. A. 2008. Trait-based community ecology of phytoplankton. Annu. Rev. Ecol. Evol. Syst. 39: 615–639.

Lunn, D. et al. 2009. The BUGS project: evolution, critique and future directions. Statistics in medicine 28: 3049–3067.

Magurran, A. E. et al. 2010. Long-term datasets in biodiversity research and monitoring: assessing change in ecological communities through time. Trends Ecol. Evol. 25: 574–582.

McGill, B. J. et al. 2006. Rebuilding community ecology from functional traits. Trends Ecol. Evol. 21: 178–185.

Michaloudi, E. et al. 2018. Reverse taxonomy applied to the Brachionus calyciflorus cryptic species complex: Morphometric analysis confirms species delimitations revealed by molecular phylogenetic analysis and allows the (re)description of four species. PLOS ONE 13: e0203168.

Monnahan, C. C. et al. 2017. Faster estimation of Bayesian models in ecology using Hamiltonian Monte Carlo. Methods in Ecology and Evolution 8: 339–348.

Novick, A. and Szilard, L. 1950. Description of the chemostat. Science 112: 715–716.

Papanikolaou, N. E. et al. 2016. Bayesian inference and model choice for Holling’s disc equation: a case study on an insect predator-prey system. Community Ecol. 17: 71–78.

Paraskevopoulou, S. et al. 2018. Differential response to heat stress among evolutionary lineages of an aquatic invertebrate species complex. Biol. Lett. in press.

Plummer, M. 2003. “JAGS: A program for analysis of Bayesian graphical models using Gibbs sampling”. In: Proceedings of the 3rd International Workshop on Distributed Statistical Computing (DSC). Ed. by K. Hornik et al.

Post, D. M. and Palkovacs, E. P. 2009. Eco-evolutionary feedbacks in community and ecosystem ecology: interactions between the ecological theatre and the evolutionary play. Philos. Trans. R. Soc. B Biol. Sci. 364: 1629–1640.

Raatz, M. et al. 2017. High food quality of prey lowers its risk of extinction. Oikos 126: 1501–1510.

Rall, B. C. and Latz, E. 2016. Analyzing pathogen suppressiveness in bioassays with natural soils using integrative maximum likelihood methods in R. PeerJ 4: e2615.

Real, L. 1977. The kinetics of functional response. American Naturalist 111: 289–300.

Reusch, T. B. H. et al. 2005. Ecosystem recovery after climatic extremes enhanced by genotypic diversity. Proc. Natl. Acad. Sci. 102: 2826–2831.

Robinson, O. J. et al. 2017. Disentangling density-dependent dynamics using full annual cycle models and Bayesian model weight updating. J. Appl. Ecol. 54: 670–678.

Rosenbaum, B. and Rall, B. C. 2018. Fitting functional responses: Direct parameter estimation by simulating differential equations. Methods in Ecology and Evolution 9: 2076–2090.

Smith, M. J. et al. 2015. Inferred support for disturbance-recovery hypothesis of North Atlantic phytoplankton blooms. J. Geophys. Res. C: Oceans 120: 7067–7090.

Stan Development Team. 2018. RStan: the R interface to Stan. R package version 2.17.3.

Taboadai, F. G. and Anadón, R. 2016. Determining the causes behind the collapse of a small pelagic fishery using Bayesian population modeling. Ecol. Appl. 26: 886–898.

Tirok, K. and Gaedke, U. 2007. Regulation of planktonic ciliate dynamics and functional composition during spring in Lake Constance. Aquat. Microb. Ecol. 49: 87–100.

Toni, T. et al. 2009. Approximate Bayesian computation scheme for parameter inference and model selection in dynamical systems. J. R. Soc. Interface 6: 187–202.

Vehtari, A. et al. 2018. loo: Efficient leave-one-out cross-validation and WAIC for Bayesian models. R package version 2.0.0.

Violle, C. et al. 2012. The return of the variance: Intraspecific variability in community ecology. Trends Ecol. Evol. 27: 244–252.

Weigelt, A. et al. 2010. The Jena Experiment: six years of data from a grassland biodiversity experiment. Ecology 91: 930–931.

Wittwer, T. et al. 2015. Long-term population dynamics of a migrant bird suggests interaction of climate change and competition with resident species. Oikos 124: 1151–1159.

